# Monocyte intrinsic NOD2 signalling inhibits pathogenic macrophage differentiation and its loss in inflammatory macrophages improves intestinal inflammation

**DOI:** 10.1101/2022.10.06.506772

**Authors:** Camille Chauvin, Daniel Alvarez Simon, Katarina Radulovic, Olivier Boulard, William Laine, Myriam Delacre, Nadine Waldschmitt, Elodie Segura, Jérome Kluza, Mathias Chamaillard, Lionel F. Poulin

**Author notes:** These authors share senior authorship (Mathias Chamaillard, Lionel F. Poulin). These authors contributed equally (Camille Chauvin, Daniel Alvarez Simon).

## Abstract

**Objective:** It is believed that intestinal recruitment of monocytes from Crohn’s Disease (CD) patients who carry NOD2 risk alleles may repeatedly give rise to recruitment of pathogenic macrophages. We investigated an alternative possibility that NOD2 may rather inhibit their differentiation from intravasating monocytes.

**Design:** The monocyte fate decision was examined by using germ-free mice, mixed bone marrow chimeras and a culture system yielding macrophages and monocyte-derived dendritic cells (mo-DCs). We next asked whether Nod2 in either monocytes or tissue macrophages have distinct resolving properties in colitis.

**Results:** Despite a similar abundance of monocytes, the intestinal frequency of mo-DCs from *Nod2*-deficient mice was lowered independently of the changes in the gut microbiota that are caused by Nod2 deficiency. Similarly, the pool of mo-DCs was poorly reconstituted with mobilized bone marrow *Nod2*-deficient cells. The use of pharmacological inhibitors revealed that activated NOD2 at an early stage of development dominantly inhibits mTOR-mediated macrophage differentiation in a TNFalpha-dependent manner. These observations were supported by the identification of a TNFalpha-dependent response to MDP that is specifically lost in CD14-expressing blood cells bearing the frameshift mutation in NOD2. Accordingly, loss of NOD2 in monocytes lowers glycolytic reserve, CD115 expression and pro-resolving features. Dietary intake of aryl hydrocarbon receptor (AHR) agonists that promotes mo-DCs generation improves colitis in *Nod2*-deficient mice to the same extent as what is observed upon macrophage ablation of Nod2.

**Conclusion:** NOD2 negatively regulates a macrophage developmental program through a feed-forward loop that could be exploited for overcoming resistance to anti-TNF therapy in CD.

**Significance of this study:** *What is already known about this subject?:*

- Loss of NOD2 function is predisposing to Crohn’s disease.
- Rapamycin, a serine/THR kinase inhibitor of mammalian target (mTOR) has been reported as potentially effective treatment in discrete subset of CD patients with refractory colitis.
- The NOD2 protein promotes the chemokine CCL2-dependent recruitment of inflammatory monocytes in response to tissue injury.
- An accumulation of CCR2-expressing monocytes and inflammatory macrophages is observed within the intestinal mucosa of CD patients including those resistant to anti-TNF therapy.
- Activated NOD2 enhances proinflammatory activity of CX3CR1^int^Ly6C^hi^ effector monocytes.
- The monocyte fate toward mo-DCs is orchestrated by the aryl hydrocarbon receptor.

*What are the new findings?:*

- NOD2 has a hierarchically dominant negative role on the mTORC-driven monocyte conversion to inflammatory macrophages independently of the changes in the gut microbiota that are caused by Nod2 deficiency.
- A defect in monocytes fate at the early stage allow the expansion of pathogenic macrophages in Nod2-deficient mice at the expense of mo-DC.
- Adoptive transfer of *Nod2*-deficient monocytes into wild-type mice was sufficient to exacerbate DSS-induced intestinal damage.
- The glycolytic reserve of monocytes and their ability to respond to M-CSF is lowered upon loss of NOD2 signalling.
- Deletion of NOD2 in macrophage improves colitis to the same extent as dietary supplementation of AHR agonist in mice.
- The recognition of the gut microbiota by NOD2 is required for *de novo* reconstitution of mo-DCs in the lamina propria of the murine intestine, while having minimal effect on the mobilization of their precursors to the intestinal mucosa.

**How might it impact on clinical practice in the foreseeable future?:** This study might contribute to the development of novel mTORC-based therapeutic strategies for improving the response to biologics by restoring the ability of circulating monocytes to reconstitute the pool of mo-DCs during homeostatic turnover and upon tissue injury. It may thereby prevent the accumulation of pathogenic macrophages in patients with loss-of-function NOD2 alleles, which fail to respond to anti-TNF and are at greater risk of developing stricturing disease.

## Introduction

The intestine is particularly enriched in phagocytes that ensure robustness in the steady state and play a role in processes of remodelling upon tissue injury. Among those, some macrophages have been seeded embryonically and self-renewed from yolk sac-derived precursor cells in mice. Under healthy conditions, intestinal macrophages have an estimated half-life of three weeks. In Human and mouse, subsets of macrophages have been characterized based on a differential expression of CD11c^1, 2^. The pool of Ly6C^low^ CX3CR1^int^ macrophages that share some features of monocyte-derived dendritic cells (mo-DCs) is then continually replenished by emigration of short-lived Ly6C^high^ monocytes from the bloodstream in mice^3^. Activation of the C-C chemokine receptor type 2 (CCR2) is a prerequisite for such leukocytes to exit from the bone-marrow in the steady state^4^. Along a continuum of differentiation stages, Ly6C^high^ monocytes may then give rise to nascent phagocytes with ascribed different functions and distinct bioenergetic programs^5^. Whereas mature macrophages are largely immotile with phagocytic capacity^6^, mo-DCs are believed to migrate for presenting protein antigens on major histocompatibility complexes class I and II (MHCI and MHCII) molecules to T cells^7-9^. The community of circulating Ly6C^high^ monocytes that are rapidly mobilized upon injury is then thought to seemingly serve as a reservoir of mo-DCs that can easily be distinguished from CD11c-expressing macrophages on the basis of ontogenetic, morphological, and gene expression criteria^10^.

In Humans, the equivalent of inflammatory monocytes are classical monocytes CD14^+^CD16^-^ that represent up to 80-95% of the large reservoir of monocytes. They can be distinguished from intermediate and non-classical subsets by their expression of well-characterized surface proteins, including CD16 (also referred as Fc gamma receptor IIIa) and the glycoprotein CD14 that acts as a co-receptor for toll-like receptor 4. While the intermediate monocytes CD14^+^CD16^+^ regulate angiogenesis and modulate effector T cell activity^11^, the nonclassical monocytes CD14^-^CD16^+^ are mobile and involved in maintenance of vascular homeostasis^12^. Recent studies demonstrated that the molecular ontogeny of human monocyte-derived cells is orchestrated by distinct transcription factors that are specifically activated by environmental cues. Comparative transcriptomic analysis revealed that the monocyte fate specification into mo-DC and monocyte-derived macrophages (mo-Mac) is at least partially coordinated by Interferon Regulatory Factor 4 (IRF4) and MAF BZIP Transcription Factor B (MAFB)^8^ respectively. Of particular importance, it has now been elegantly demonstrated that the monocyte fate toward mo-DCs is orchestrated by the aryl hydrocarbon receptor^8^, defect of which is associated with the susceptibility of several common diseases, including Crohn’s disease (CD)^13^. Likewise, monocytes may give rise to mo-DCs upon inhibition of the mammalian target of rapamycin (mTOR) pathway^14^ for which the sustained activation in CD is likely a consequence of a genetically predisposed defect in autophagy^15^. Interestingly, an increased number of inflammatory macrophages is observed within the intestinal mucosa of CD patients at the expense of their proresolving counterpart^16^. Those data were supported by recent single-cell analysis of inflamed tissues from CD, which revealed the presence of discrete subset of pathogenic macrophages within the diseased intestine of CD patients that fail to respond to anti-TNF therapy^17^. It is thereby tempting to speculate that a defect in development of monocyte-derived phagocytes may allow the expansion of pathogenic macrophages that maintain a TH1-biased CD4 T cell response through the production of inflammatory and fibrogenic effectors.

Genetic variants in the *NOD2* gene confer an increased susceptibility to CD, likely due to loss of NOD2 function. Furthermore, patients who carry NOD2 risk alleles are at greater risk of developing stricturing disease^17^. The Nucleotide binding Oligomerization Domain (NOD)-like receptor NOD2 is a cytosolic sensor of bacterial muramyl dipeptide (MDP). MDP is an active component in Freund’s complete adjuvant and derivatives have been synthesized for improving their pharmacological properties. Intact bacteria on already differentiated monocyte-derived DCs demonstrate a role of MDP on autophagy induction, bacterial destruction and antigen presentation ^18^. Furthermore, probiotics induce an anti-inflammatory effect on mo-DC in a Nod2- and strain-specific manner^19^. Despite substantial efforts that were made in studying how the homeostatic trafficking of monocytes is controlled by NOD2, it remains unclear whether NOD2 may orchestrate their differentiation into developmentally distinct subset of cells that are specialized for maintenance of immune surveillance. After stimulating their exit from the bone marrow, classical monocytes have the capacity of being converted into non-classical cells in a NOD2-dependent manner^20^. Besides this phenomenon, it is now well established that sensing of bacterial endotoxin promotes the mobilization of inflammatory monocytes, which can develop into cells with a typical probing morphology and with critical features of mo-DCs including cross-priming capacities of cell-associated antigen to CD8^+^ T cells^7^. It suggests the likelihood that loss of NOD2 may directly inhibit the development of programmable cells of monocytic origin into inflammatory macrophages. While mo-DCs are markedly less abundant within the healthy intestine than macrophages, it does not exclude the possibility that mo-DCs may play an essential role in intestinal homeostasis. One may then consider that NOD2 may influence the differentiation of bone marrow precursors into tissue phagocytes in a context-dependent manner.

In this study, we provide experimental evidence that NOD2-dependent bacterial sensing by programmable monocytes inhibits their differentiation in macrophages at early stage of their development. In agreement, ablation of Nod2 in terminally differentiated macrophages had no influence on the frequency of nascent macrophages. Such a phenomenon occurs even when their development from circulating monocytes into primitive macrophages is enforced upon either injury or activation of the metabolic signalling node mTORC1 that controls terminal differentiation of myeloid progenitors^21^. This mutually exclusive switch on early steps of phenotypic developmental stages relied on the glycolytic-mediated control of monocyte responsiveness to M-CSF by the bacterial sensor NOD2. We demonstrated that recognition of the gut microbiota by NOD2 is required for *de novo* reconstitution of mo-DCs that occupy the lamina propria of the murine intestine, while having minimal effect on the mobilization of their precursors to the intestinal mucosa. In agreement with such effect of NOD2 at early stage on the monocyte fate, we found that ablation of Nod2 in terminally differentiated macrophages improves colitis, while having no influence on the frequency of nascent phagocytes when populating the intestinal niche at either steady state or upon injury. Given that MDP is physiologically present in high concentrations within the intestinal lumen, our study set the stage to modulate NOD2-dependent signalling at the early M-CSF-dependent monocytic stage to avoid the activation of default developmental pathways, including mTORC1. These alternative developmental switches are probably leading to anti-TNF failure and stricturing complications through accumulation of inflammatory macrophages in the intestine of CD patients.

## Results

### Recognition of the gut microbiota by Nod2 regulates reconstitution of intestinal mo-DCs from mobilized monocytes

To exclude the potential influence of opportunistic pathobionts that may have been present in *Nod2*-deficient mice, the faecal microbiota from wild-type mice was transplanted in either germ-free recipients that are deficient or not for NOD2 (SFig1A). The composition of CD11c+MHCII+ and CD11c-MHCII+ subsets of mononuclear phagocytes was analysed four weeks after colonization for making their gut microbiota similar to the one of control mice (SFig1B). The frequency and absolute numbers of CD11c+MHCII+ and CD11c-MHCII+ subsets of mononuclear phagocytes were not significantly different in the ex-germ free mice exposed to WT microbiota (SFig1B). These results suggested that changes in the composition of the gut microbiota that are pre-existing in the *Nod2*-deficient mice could not be sufficient for influencing the frequency of mononuclear phagocytes within the colonic lamina propria in mice. To investigate more precisely whether the gut microbiota may impact the mononuclear phagocyte composition of the colonic lamina propria in a Nod2-dependent manner, live single cell suspensions were prepared from the colon of specific pathogen-free (SPF) and germ-free (GF) mice before being analysed by multiparameter flow cytometry (Fig1A). After excluding dead cells, doublets and lineage-positive cells, a CD11c-CD11b dot plot was subdivided according to the subset of hematopoietic cells expressing CCR2, Ly6C and further subdivided on the absence or presence of major histocompatibility complex class II (MHCII) at their cell surface (Fig1A). The frequency of activated MHCII^+^ monocytes is reduced within the lamina propria of the colon of GF mice when compared to SPF condition (Fig1A,B and SFig2). Accordingly and as observed in a series of studies^3, 22^, the proportion of mature tissue macrophages was heightened in the colon of SPF mice when compared to GF condition (Fig1A,C and SFig2). Those differences were obviously maintained even in *Nod2*-deficient mice. By contrast, the expression level of MHCII on the classical subset of monocytes was similar within the colon of SPF *Nod2*-deficient mice as what was observed in GF condition (Fig1B and SFig2). As a comparison, we next studied in greater details the heterogeneity within the CD11c-expressing cells. As NOD2 has been shown to shape the recruitment of CD103^+^ DCs in the gut lamina propria^23^, we reasoned that the loss of Nod2 signalling may have impaired either the trafficking or the development of some discrete subsets of conventional dendritic cells that do not express Ly6C and are largely dependent on GM-CSF, also known as colony-stimulating factor 2 or Csf2. To our surprise, only minor fluctuations were detected in the absolute numbers of cDC1 and cDC2 that co-express or not CD11b respectively (Fig1D). Interestingly, we observed that the colon of SPF and GF *Nod2*-deficient mice harboured a lower frequency of cells co-expressing the surface molecules CD11b, CD11c, Ly6C, CCR2 and MHCII, which is a subset commonly referred to as mo-DCs^24^, as compared to SPF WT mice (Fig1C). As this difference is lost in GF mice, these results suggested to us that NOD2-dependent differences in frequency of mo-DCs could result from a cell-intrinsic mechanism instead of being microbiota dependent. Given that the gut microbiota does not regulate the abundance of cDC1, the decreased abundance of mo-DCs was not likely due to a competition with cDC1 to occupy the colonic niche under homeostatic conditions. This said, the number of CD11c-expressing macrophages was too low to conclusively apprehend potential differences with our experimental setting. Altogether, our data indicate that the Nod2-dependent recognition of the gut microbiota by monocytes when entering the colon from the blood is an important means by which NOD2 could prevent the replenishment of macrophages from mobilized monocytes and subsequently facilitate on-demand accumulation of intestinal mo-DCs.

**Figure 1.**
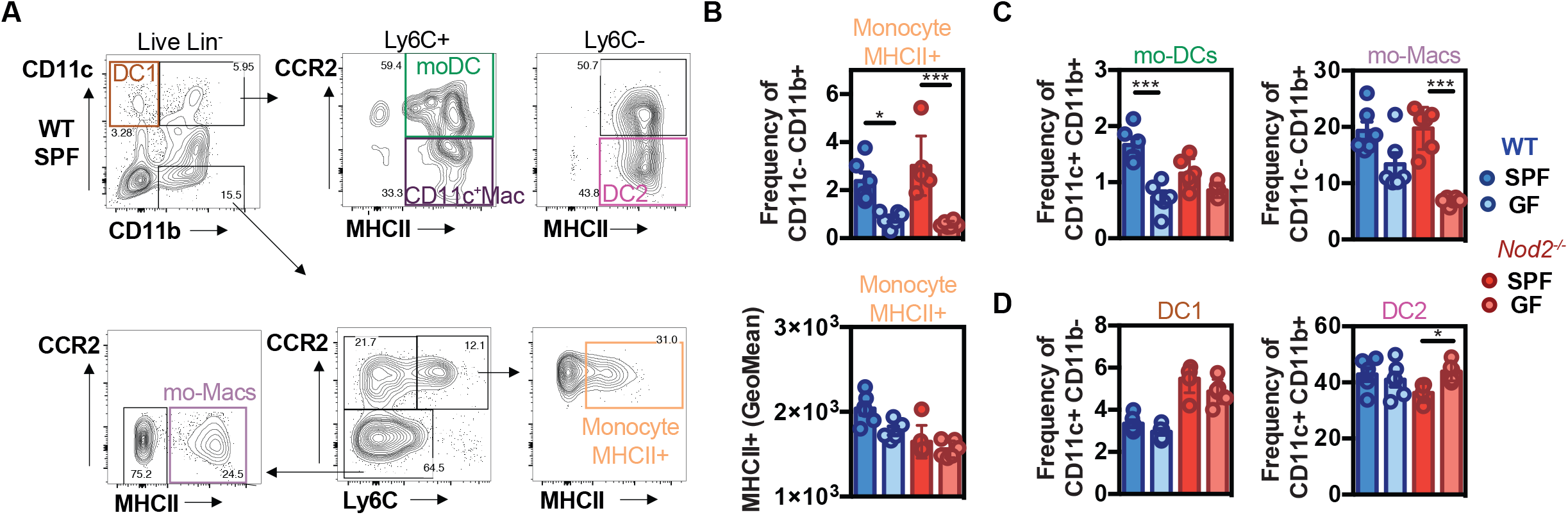
Default of conventional DCs and monocytes-derived cells recruitment in NOD2-/- mice under microbiota deprivation. Colon Lamina Propria Mononuclear Cells (LPMC) frequency analysis in Wild-type (WT) (blue) and NOD2-/- (red) mice raised under Specific-Pathogen Free (SPF)(sharp color) and Germ-Free(GF)(light color) conditions. (A) Gating strategy to determine the frequency of conventional DC1 (CD11c+CD11b-), of mo-DCs (CD11c+CD11b+Ly6C+CCR2+MHCII+), of CD11c+ Macs ((CD11c+CD11b+Ly6C+CCR2-MHCII+) of conventional DC2 (CD11c+CD11b+Ly6C-CCR2-MHCII+), and the frequencies of mo-Macs (CD11c-CD11b+Ly6C-CCR2-MHCII+), and Mo-MHCII+ (CD11c-CD11b+Ly6C+CCR2+MHCII+), based on their CCR2, Ly6C and MHC levels, after exclusion of Lineage and doublet cells. (B) The frequency and MHCII GeoMean of CCR2+Ly6C+MHCII+ activated monocytes was evaluated in the monocyte-derived cells CD11c-CD11b+. (C) The frequency of mo-DCs was evaluated in the Ly6C+ cells. The frequency of CCR2-Ly6C-mo-Macs was evaluated in the CD11c-CD11b+ monocyte-derived cells while. (D) Frequency of DC1 cells (CD11c+CD11b-gate) and DC2 cells (Ly6C-gate). Bars indicate mean ± SEM (four to six mice per group). Statistical significance was assessed by two-way ANOVA. *, P<0.05; **, P<0.01.

### Lack of Nod2 results in a competitive disadvantage for the mo-DCs pool within the colon and the peritoneal cavity at steady-state in mice

Since a delayed appearance of mo-DCs is observed within the intestine of SPF *Nod2*-deficient mice at steady state, we next determined whether this phenomenon is a consequence of an impaired mobilization of monocytes from the bone marrow as it was observed in C3H/HeJ Tlr4 mutant mice^7^. Alternatively, maturation of Ly6C^high^ monocytes can follow different paths such a mo-DCs or CD11c^+^ macrophages which are considered as an intermediate between monocytes and macrophages^1^. If Nod2 is intrinsically required for the development of Ly6C^high^ monocytes into mo-DCs in mice, inappropriate conversion of Ly6C^high^ monocytes into mo-DCs would be expected to promote accumulation of macrophages. To this end, we realized mixed bone marrow chimera mice. *Nod2*-deficient animals were lethally irradiated, and 24h later reconstituted with equal amounts (eg. 50:50) of bone marrow cells from WT (CD45.1) and *Nod2*-deficient mice (CD45.2) (Fig2A). Peritoneal content of mo-DCs and mo-Macs within the peritoneum^8^ and colonic tissue was analysed 8 weeks after bone marrow reconstitution as described previously (Fig2). Within the peritoneum of *Nod2*-deficient competitive bone marrow chimeras, we observed a lowered relative proportion of mo-DCs when compared to mo-Macs (Fig2B,C). Similar results were obtained in their colon in which the number of macrophages was significantly heightened (SFig3). We next assessed whether the capacity of blood monocytes to be recruited into the colon may depend on Nod2-mediated recognition of the gut microbiota. Under these experimental conditions, monocytes were similarly recruited in the colon despite the loss of Nod2 signalling in radioresistant cells (Fig2D). We next defined the extent to which inflammation may differentially alter the proportion of bone marrow-derived phagocytes in the colon of *Nod2*-deficient chimeras when compared to that in similarly treated control chimeras. Dextran sodium sulfate (DSS) was then administered in the drinking water of mixed-bone marrow chimera mice to induce acute colitis (Fig2E-G). Such an established preclinical model of colitis is characterized by epithelial erosion, crypt loss, ulceration and infiltration of immune cells. The evaluation by flow cytometry revealed a similar reconstitution of Ly6C^high^ monocytes in the blood, and in the colon, before and after DSS injury regardless of the genotype of donor cells chimeras (Fig2E, SFig4A,B). In agreement, the ratio of colonic and blood monocytes was equivalent between each donor cell (Fig2F). Likewise, the number of colonic Ly6C^high^ monocytes that are activated or not were not affected by Nod2 deficiency in response to DSS (SFig4C). Furthermore, no differences in the body weight loss was noticed at the time of autopsy (data not shown). Upon injury, the number of conventional dendritic cells and the relative proportion of mo-DCs compared to mo-Macs were equivalent in the colon of mixed bone marrow chimera mice regardless of the genotype from donor bone marrow cells (Fig2G). Although the reconstitution of Ly6C^high^ monocytes in the blood was not affected by Nod2 deficiency, CD11c-expressing macrophages that are deficient for Nod2 were present in a greater number in comparison to WT donor cells (Fig2G). Overall, these results support the possibility that under inflammatory conditions the generation of colonic macrophages is likely inhibited by Nod2 signalling in tissue Ly6C^high^ monocytes.

**Figure 2.**
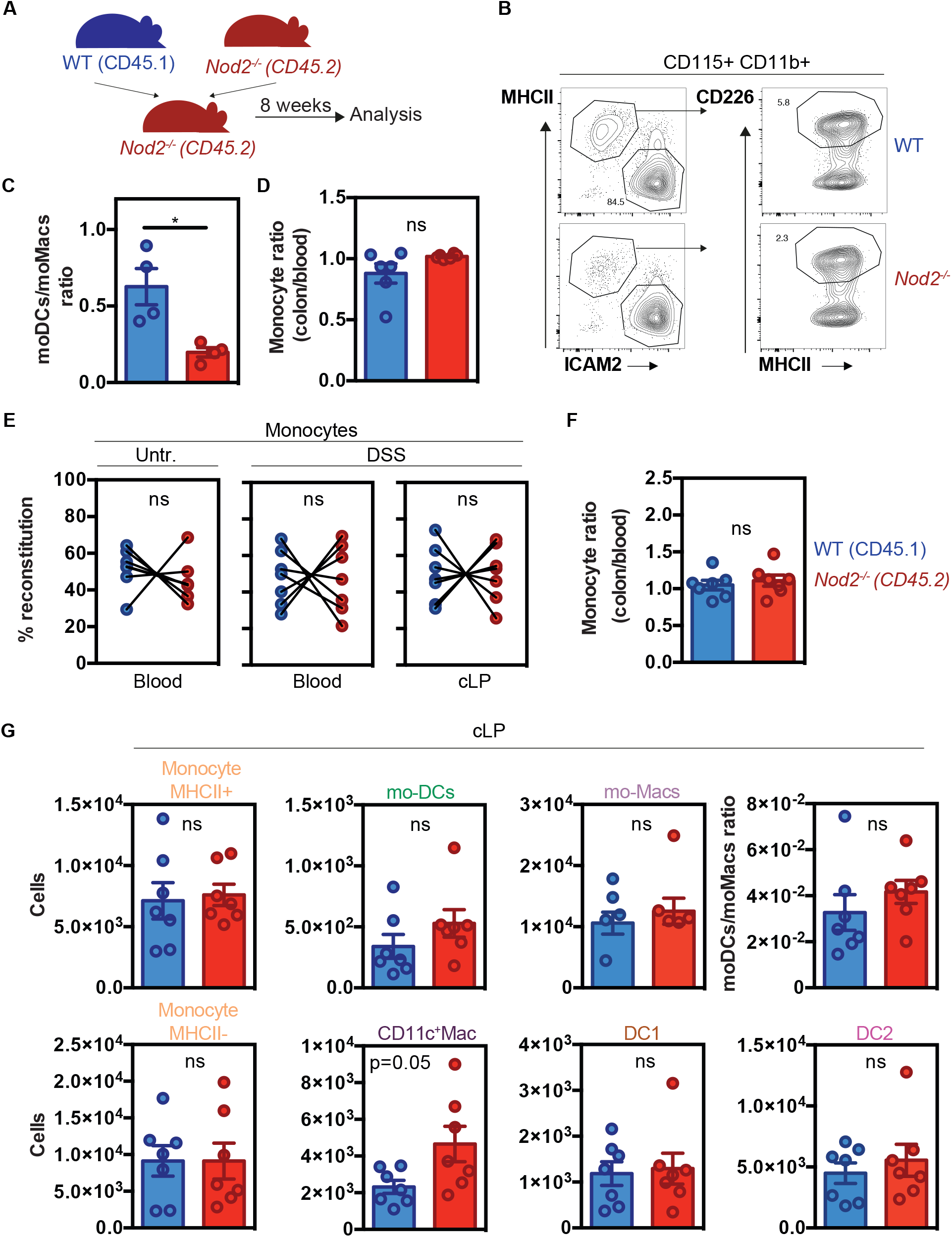
Nod2 signalling is required in bone marrow cells to reconstitute the mo-DCs pool in the peritoneal cavity despite similar monocyte recruitment. (A) Experimental protocol. Mixed bone marrow chimeras were generated by transferring WT (expressing CD45.1)(blue) and Nod2-/- (expressing CD45.2)(red) cells in a 1:1 ratio into lethally irradiated Nod2-deficient recipients. Cells were isolated from the blood and the peritoneum 8 weeks after reconstitution. (B) Peritoneal cells were harvested and the proportion of Mo-Mac (MHCII-ICAM2+) and mo-DCs (MHCII+CD226+) was determined by flow cytometry by gating within the CD115+CD11b+ cells. (C) Ratio of mo-DCs/mo-Macs from frequencies obtained in B (4 mice). (D) Reconstitution index of Ly6C+ monocytes in the colon (CD11c-CD11b+Ly6C+CCR2+ gate) divided by the same ratio in the blood (6 mice). (E) 8 weeks after reconstitution, mixed chimera mice were treated with a 5-day course of 2% DSS (7 mice)(E-G). (E) Monocytes content in the blood of untreated and blood and colonic lamina propria cells (cLP) in DSS-treated mice. (F) Ratio of WT and Nod2-/- monocyte frequencies (colon/blood). (G) Colonic cells were analyzed as described in Figure 1. Numbers of mono MHCII+, mono MHCII-, mo-DCs, CD11c+Macs, mo-Macs, DC1 and DC2 in the cLP (number of cells per million of live cells). Ratio of mo-DCs/mo-Mac from total cell number. Bars indicate mean ± SEM. Statistical significance was assessed by non-parametric Mann-Whitney U test. * P<0.05.

### Nod2-deficient monocytes aggravated DSS-induced colitis

Since NOD2 activation is required for the conversion of Ly6C^high^ into Ly6C^low^ monocytes^20^, we adoptively transferred Ly6C^high^ monocytes isolated from the bone marrow from *Nod2*^-/-^ into WT mice at 4.5 days after the start of a 8-day course of 2% DSS. This sulfated polysaccharide is known to rapidly recruit circulating monocytes into the inflamed colonic lamina propria^25^. Interestingly, WT mice that received *Nod2*^-/-^ Ly6C^high^ monocytes had about ∼15% decrease of the initial body weight, as early as day 8 after the start of DSS treatment when compared to mice that received Ly6C^high^ monocytes from WT animals (blue circle) (Fig3A). In line with an enhanced body weight loss, WT mice injected with *Nod2*^-/-^ Ly6C^hi^ monocytes also had a slight decreased colon length at this timepoint (Fig3B), despite no changes in the histological score (Fig3C-D). Accordingly, injection of Nod2^-/-^ Ly6C^hi^ monocytes in WT mice resulted in a significant decrease in colitis-associated transcripts levels such as *Ido-1* and *Ifng* (Fig3E). This is in agreement with the previously reported decreased expression of *Ido-1* and *Ifng* in another model of colitis^26^. In contrast, we did not observe a significant decrease in transcript level of *Il17a* (p=0.06)(Fig3E), *Tnfa, Il1b, Il6, Spp1, Mmp13* and *Ptgs2* into the colon of these mice when compared to PBS-injected mice (data not shown). Thus, Nod2 activity in Ly6C^hi^ inflammatory monocytes plays an important role in limiting DSS-induced damage.

**Figure 3.**
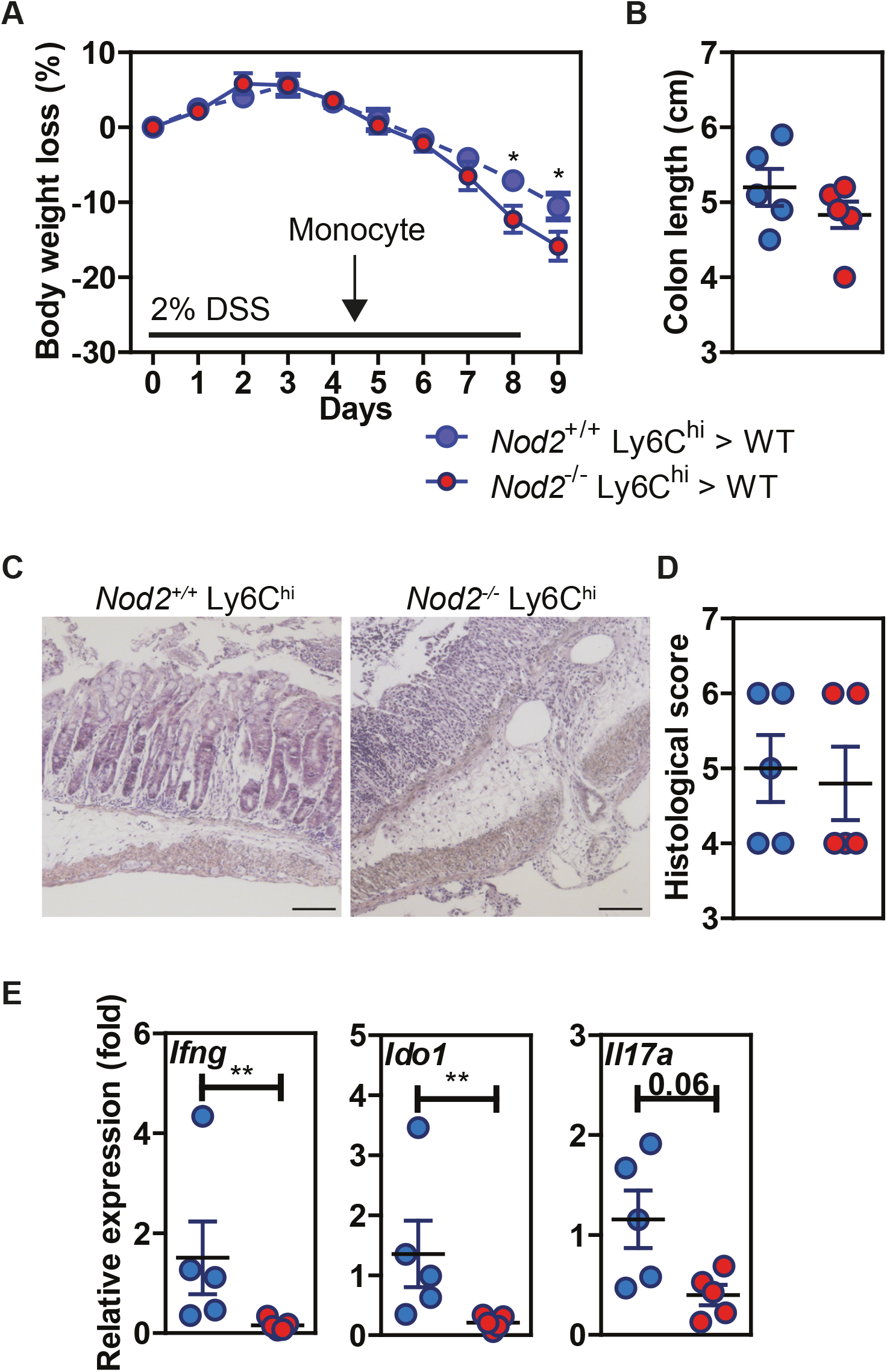
Nod2-deficient monocytes aggravated DSS-induced colitis. Nod2+/+ (blue circle) and Nod2-/- (red circle) Ly6Chi monocytes were transferred into WT mice 4.5 days after the start of a 8-day course of 2% DSS. (A) The weight loss was monitored during DSS-mediated colitis, as well as the colon length at day 9 (B). (C) Histological sections in H&E were done to confirm tissue damages. (D) Histological score determined at the sacrifice. (E) RT-qPCR on colonic tissues were done to measure *Ifng, Ido1, Il17a*. Data are representative of 2 experiments with five mice per group. Bars indicate mean ± SEM. Statistical significance was assessed by non-parametric Mann-Whitney test. **, P<0.01 were considered statistically significant.

### NOD2 signalling enhances the yield of mo-DCs from monocytes by inhibiting their differentiation in macrophages at an early stage of development

In order to establish whether activation of Nod2 signalling may intrinsically inhibit the differentiation of monocytes that are recruited on demand into macrophages, MDP was added at the start of the culture of mouse bone marrow cells with the conditioning media from J558 cells that continuously secrete GM-CSF. Culture of mouse bone marrow cells with the growth factor GM-CSF is a widely used protocol to generate either mo-Macs or mo-DCs within the CD11c^+^MHCII^+^ fraction (Fig4A)^27^. The percentage of CD115+CD64+ mo-Macs was significantly diminished when bone marrow cells were cultured with MDP as compared with medium alone (Fig4B-C). Interestingly, the output of the culture with MDP significantly increased the frequency of mo-DCs (Fig4B-C). We then verified if similar results could be observed in *in vitro* culture of human monocytes. We therefore used a well-established cocktail of conditioning soluble factors to generate yields of mo-DCs from human peripheral blood CD14^+^ monocytes^28,29^. In our hands, it has been appreciated that both mo-DCs and mo-Macs may arise from human monocytes when being cultured with the addition of GM-CSF and interleukin-4 (IL-4)^30^. Light microscopy revealed that monocytes acquired a less round shape as early as 24 hours after being exposed to MDP (Fig4D). After 5 days of stimulation, we noticed the formation of a homotypic cluster of adherent cells with a probing morphology (Fig4D). To support this, both adherent and non-adherent cells were harvested and stained at their surface for CD14, CD16, CD1a, and HLA-DR. Live and singlet cells were gated on HLA-DR-expressing cells and were systematically analysed on day 6 (Fig4E-G). As what was observed in culture with M-CSF, Tumor Necrosis Factor-alpha (TNF-a) and IL-4^31^, early presence of MDP into such a culture system reduced the frequency of mo-Macs by fivefold (Fig4F), while addition of LPS drastically diminished the frequency of mo-DCs in favour of mo-Macs that are expressing higher levels of CD16 (Fig4E-G, lower part). Conversely, MDP treatment at the early stage of the culture efficiently enhanced the incidence of mo-DCs that express higher levels of CD1a, which has long been known as a marker of *in vitro* generated mo-DCs (Fig4G, upper part). By contrast, the yield of mo-Macs was similar between GM-CSF and IL-4 culture of monocytes that are treated or not with MDP for the last two days of culture (data not shown). This is in agreement with the time-restricted ability of soluble factors such as IL-4 or TNF-alpha (TNF-α) to induce monocyte differentiation process towards mo-DCs in the first 72h of development ^32, 33^. This result is consistent with the steadily decline of NOD2 expression during the first 72h of the monocyte culture in the presence of GM-CSF and IL-4^33^. Accordingly, qRT-PCR analysis revealed a lowered expression of NOD2 in terminally differentiated mo-DCs when compared to naïve monocytes (SFig5), which is suggesting that the expression of NOD2 is temporally regulated during monocytes differentiation. These results suggest that early NOD2 signalling conditions the differentiation of nascent phagocytes into cells that promote inflammatory responses.

**Figure 4.**
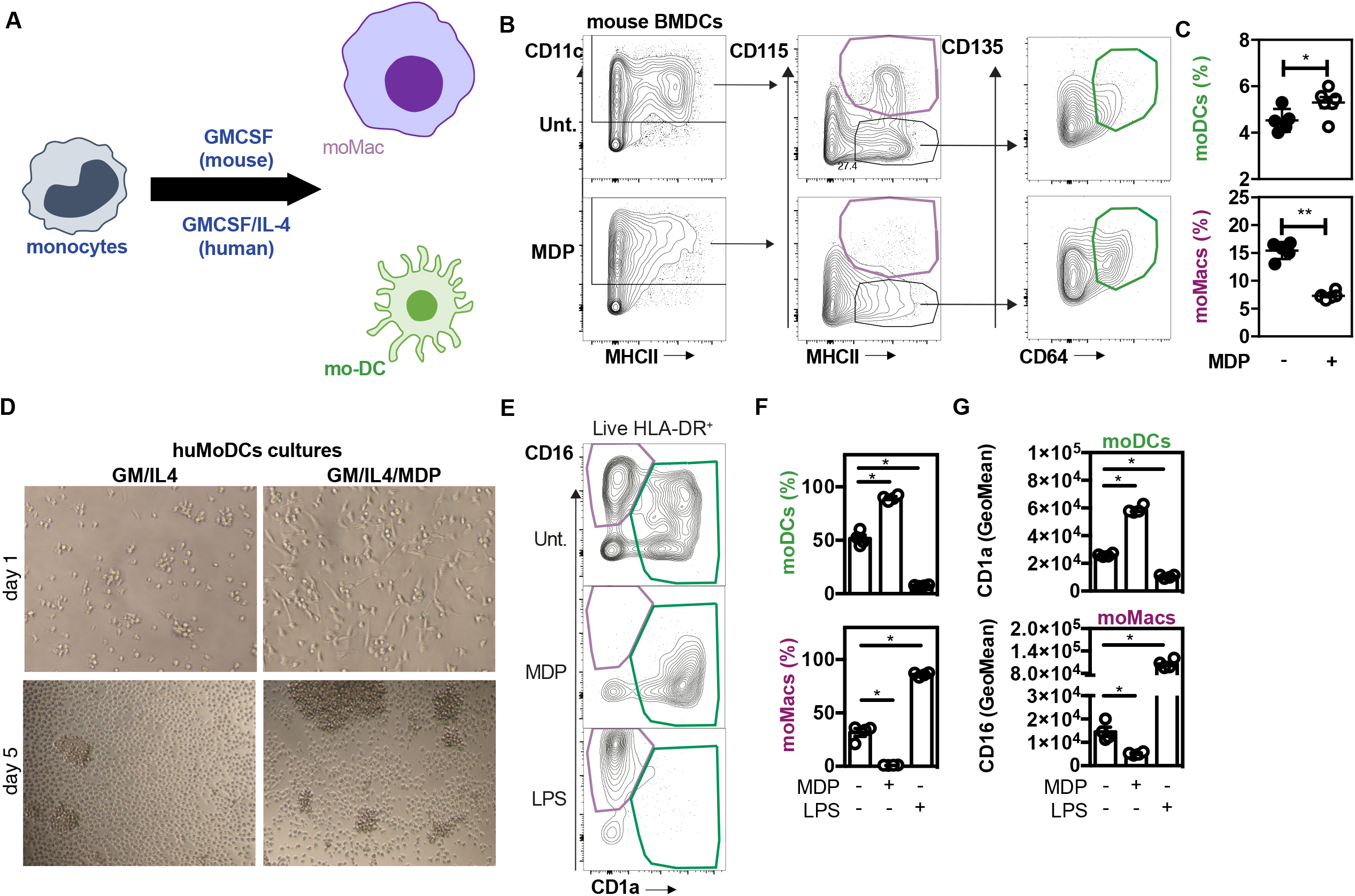
Nod2 stimulation is responsible for a mo-Macs/mo-DCs switch. (A) Experimental protocol. (B) Gating strategy for flow cytometry analysis of BMDCs upon MDP treatment. Bone-marrow derived cells were in vitro generated for 7 days in GM-CSF in the presence or not of MDP (10ug/ml). Mo-DCs were gated as CD11c+ MHCII+ CD115-CD135+ CD64+ (green) and mo-Macs were gated as CD11c+ MHCII+ CD115+ (purple). Frequencies are calculated from the CD11c+ gate. (C) Frequencies of mo-DCs and mo-Macs in the mouse BMDC culture stimulated or not with MDP at the start of the culture. (D) mo-DCs were generated by culturing CD14+ circulating human monocytes with GM-CSF and IL-4. MDP was added at the beginning of the culture. Morphology of the differentiating cells at day 1 and day 5 of culture in the presence or not of MDP. (E) The impact of MDP or LPS stimulation at the start of the culture on mo-Macs (CD16+CD1a-) and mo-DCs (CD16-CD1a+) differentiation was evaluated by flow cytometry at day 6, after gating on live HLA-DR+ cells. (F) mo-DCs and mo-Macs frequencies. (G) CD1a and CD16 fluorescent mean intensity (GeoMeans) (n=5). These data are representative of at least 4 independent experiments with different mice (B-C) or donors (D-G). Bars indicate mean ± SEM. Statistical significance was assessed by non-parametric Mann-Whitney test. *, P<0.05; **, P<0.01.

### Loss of Nod2 expression in tissue macrophages and restoration of mo-DC differentiation by dietary AHR agonist improve colitis recovery

To determine whether Nod2 expression in terminally differentiated macrophages is intrinsically required for the reconstitution of mo-DC at steady state and upon injury, Nod2^fl/fl^ mice were crossed with mice expressing the recombinant Cre in myeloid cells^34^. As expected, the expression of NOD2 was lowered in the peritoneal macrophages (but not on CD4 T cells) from those animals, that are referred to as Nod2ΔLyzM, as compared to their Nod2^fl/fl^ littermates (SFig6). At steady state, flow cytometry analysis of mononuclear phagocytes from the large intestine of littermates revealed that the acquired loss of NOD2 in terminally differentiated macrophages was not sufficient to trigger significant fluctuations of each discrete subsets of phagocytes as what observed in the intestine of *Nod2*-deficient mice (Fig5). Upon injury, an increased proportion of mo-DCs coincided with a drastic decrease of mo-Macs within the diseased colon from either Nod2^fl/fl^ or Nod2ΔLyzM littermates. Given that LysM is expressed at a later stage of monocytopoiesis, these findings are in agreement with a function of NOD2 on the early conversion of monocytes into mo-DCs. To further study the role of NOD2 signalling on the function of terminally differentiated macrophages, THP1 cells were cultivated with Phorbol 12-myristate 13-acetate (PMA-Mac) for 2 days followed by 3 days of rest, as described previously ^35^. PMA is a high affinity activator of protein kinase C. Specifically, the secretion of inflammatory cytokines was measured when treating PMA-Mac with LPS during 24h, followed by 24h treatment with MDP. In agreement with the aforementioned data using Nod2ΔLyzM, a greater responsiveness to subsequent MDP stimulation was noticed when PMA-Mac were primed with LPS as determined by quantification of TNF-α (data not shown). These results suggested that terminally differentiated macrophages may undergo a phenotypic switch to a proinflammatory function upon activation of NOD2 signalling. We next assessed whether NOD2 expression in macrophages is required for appropriate healing of injured tissue to occur. Daily monitoring of body weight and signs of colitis during the DSS colitis model revealed that Nod2ΔLyzM were losing less weight than control littermates (Fig5B). As early as seven days after DSS treatment, the colon shortening was decreased in Nod2ΔLyzM mice (Fig5C-D), which is coherent with a less damaged colon assessed by histological sections (Fig5E-F). Along the same lines, inflammatory markers such as *Il-6* and *Spp1* were less expressed as compared to control animals (Fig5G). However, we did not notice any changes in the *Tnf-α* expression. We also noticed lowered spleen weight when compared to control littermates (Fig5H). Consequently, those findings suggested that early Nod2 signalling may progressively promote the differentiation of rapidly mobilized monocytes into ontogenetically related cells with specific features of mo-DCs by inhibiting their conversion into mo-Macs within the lamina propria^36^. These results led us to assess whether the lowered mo-DC content in the colon of *Nod2*-deficient mice could be associated with, or even be responsible for, their greater susceptibility to colitis. To this end, the availability of AhR ligand was artificially enhanced by feeding DSS-exposed mice with an experimental diet enriched for indole-3-carbinole (I3C) that promotes mo-DCs generation^8^. The body weight loss that was monitored in mutant mice was improved upon dietary intake of aryl hydrocarbon receptor (AHR) agonists when compared to animals who received a control diet (SFig7). Overall, our data, MDP sensing by monocytes could promote their early conversion into mo-DCs for maintenance of intestinal homeostasis at the expense of inflammatory macrophages.

**Figure 5.**
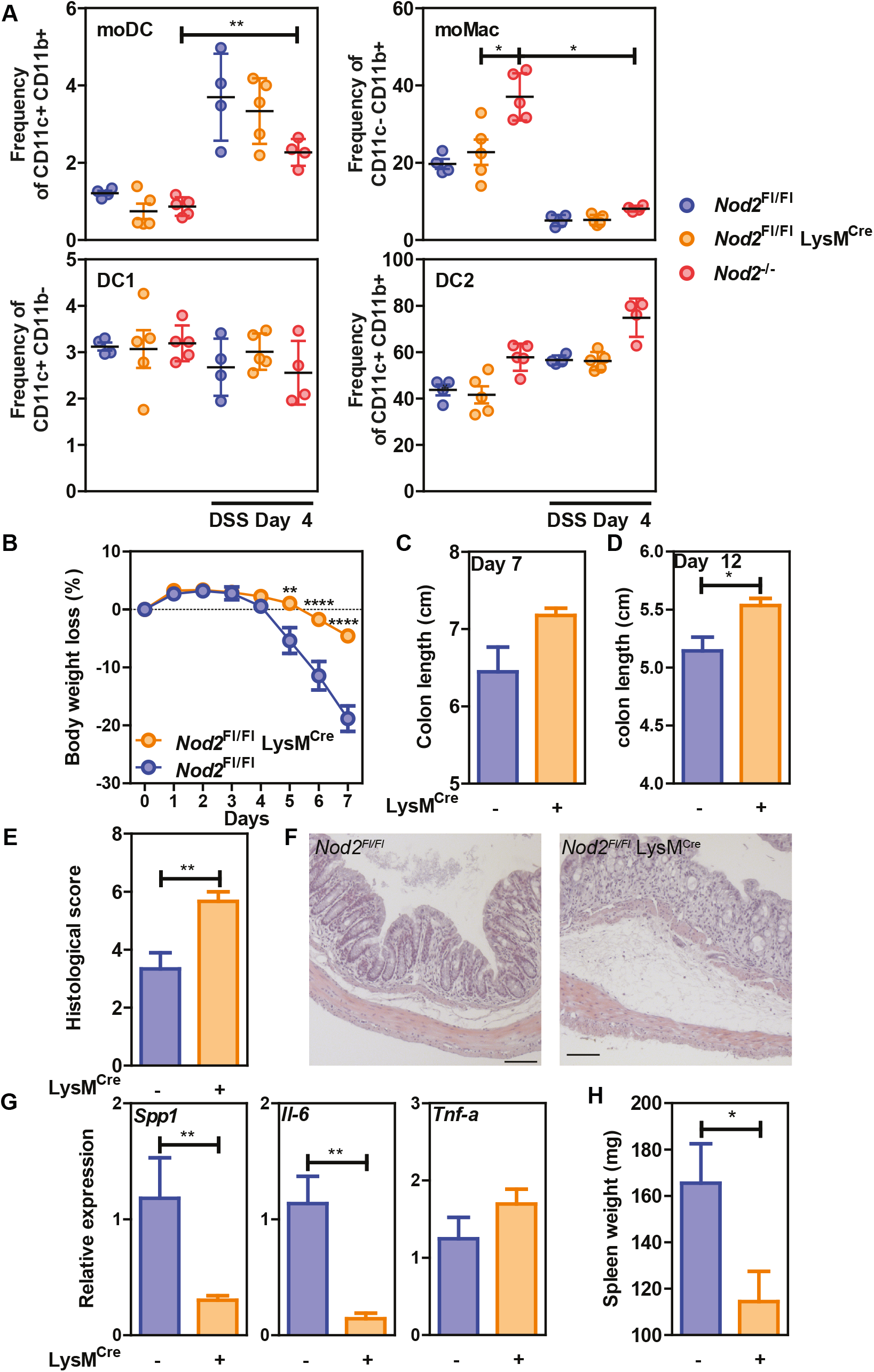
Nod2 deficiency in mature mononuclear phagocytes is protective in colitis. (A) Nod2ΔLyzM were compared to control flox mice and Nod2-/- after induction of DSS-mediated colitis. The frequencies of mo-DCs, mo-Macs, DC1 and DC2 were evaluated in the colon before and after DSS induction, as described in Figure 1 (4-5 mice per group). (B) The weight loss was monitored during DSS-mediated colitis, as well as the colon length at day 7 (C) and day 12 (D) and the spleen weight (H). Histological sections in H&E were done to confirm tissue damages (E-F). Representative of 2 independent experiments. (G) RT-qPCR on colonic tissues were done to measure *Il-6, Tnf-a*, and *Spp1* at day 7 after DSS induction (5-7 mice per group). Bars indicate mean ± SEM. Statistical significance was assessed by non-parametric Mann-Whitney test, or by two-way ANOVA, Bonferroni’s multiple comparisons test (B). *, P<0.05; **, P<0.01; ****, P<0.001.

### NOD2 signalling acts downstream of the mTORC1 pathway for licensing early stage bifurcation of monocytes commitment

We next asked how Nod2 signalling in monocytes may modulate the unique property of monocytes to differentiate into macrophages that are phenotypically and functionally different from mo-DCs. To this end, we investigated the signalling events after Nod2 stimulation with a focus on the components of the mechanistic target of rapamycin (mTOR) signalling pathway that are both involved in the generation and activity of tissue resident peritoneal macrophages *in vivo*^*37,38*^. In order to assess the importance of NOD2 signalling on these signalling pathways under MDP stimulation, we made use of the Human monocytic cell line THP1 that express NOD2^39^. This cell line has been extensively used to study the development and function of human monocyte-derived phagocytes. We first asked whether MDP stimulation of monocytes might fail to differentiate into mo-DCs after enforced activation of mTORC1. Specifically, we used MHY1485 which is a cell permeable activator that targets the ATP domain of mTORC1 but not of mTORC2. In agreement with our hypothesis, adding MHY1485 to monocytes at the start of the culture with GM-CSF and IL-4 enhanced the proportion of macrophages (Fig6A,B) to the same extent as what was observed with GM-CSF culture of mouse myeloid progenitors^40^. Interestingly, MDP addition led to a significant decrease of macrophage frequency even during the MHY1485 treatment suggesting that NOD2 may act downstream of mTORC1 during monocyte differentiation (Fig6A,B). We then reasoned that inhibition of the mTOR pathway could prevent the conversion of monocytes into macrophages to the same extent as what was observed in response to MDP. In line with our hypothesis, the proportion of mo-DCs was enhanced upon treatment of monocytes with wortmannin that acts upstream of the mTOR pathway to the same extent as what observed with MDP. Wortmannin is a non-specific, covalent inhibitor of phosphoinositide 3-kinases (PI3K) that is also used for suppressing autophagy by interfering with autophagosome formation. Cell toxicity was avoided as much as possible by treating cells for only 24 hours (data not shown). Similar results were obtained with the prototypic mTORC1 inhibitor referred to as rapamycin that is able to induce autophagy by potentiating LC3 lipidation (Fig6A,B). These data are in agreement with the greater proportion of mo-DCs that is observed upon treatment with the mTORC1 inhibitor temsirolimus^14^. These results suggested us that NOD2 activation might negatively regulate the ability of the PI3K pathway to license metabolic reprogramming of monocytes at an early stage of development.

**Figure 6.**
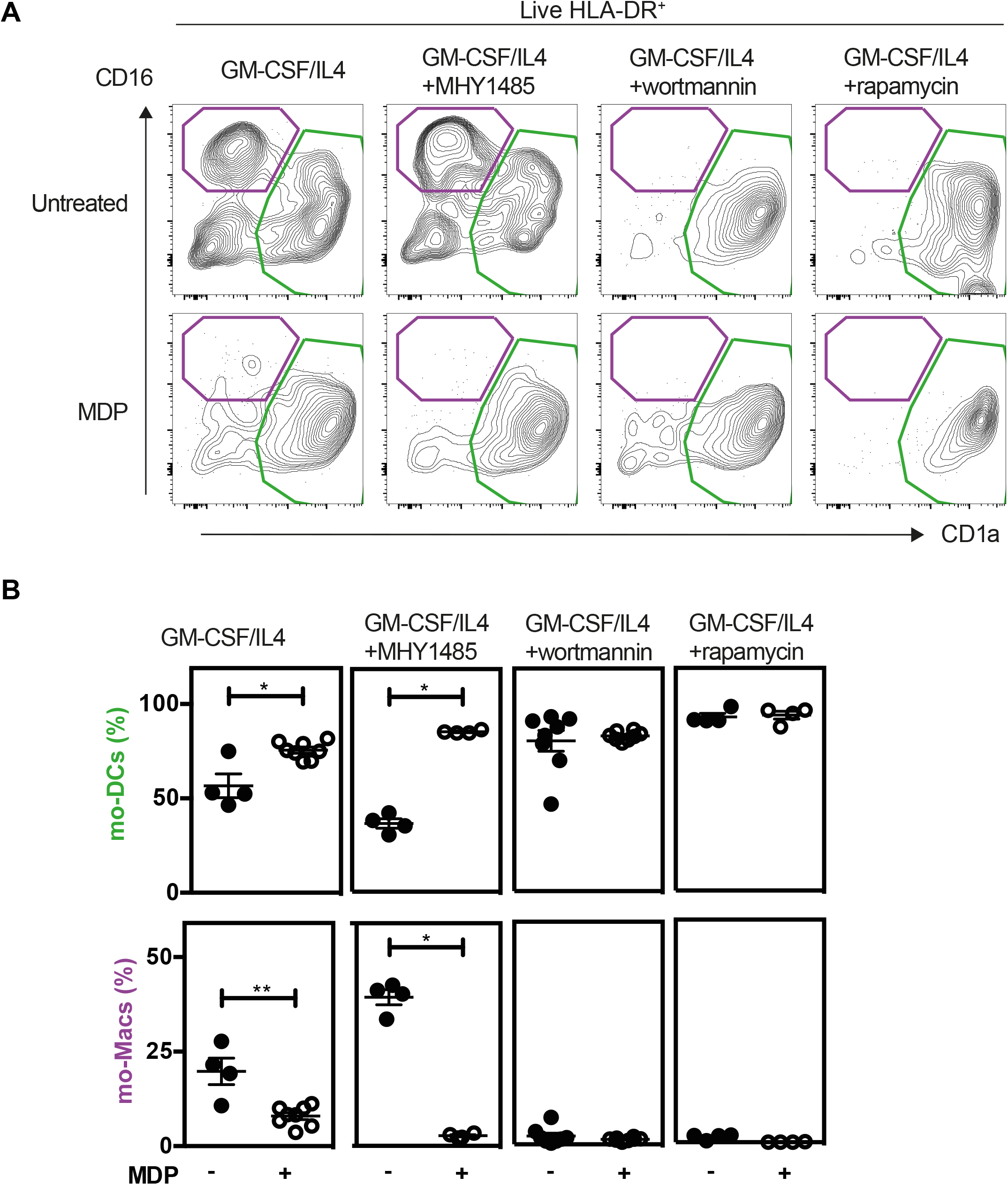
MDP enhances the differentiation of Mo-DCs in a mTORC1 independent manner. Monocytes were treated or not with MDP, the mTOR activator MHY1485, the PI3K inhibitor wortmannin or the mTOR inhibitor Rapamycin. Mo-DCs (CD1a+) and mo-Macs (CD16+) frequencies were assessed by flow cytometry using the HLA-DR, CD16 and CD1a markers. (A) Contour representing the mo-DCs (green) and mo-Macs (purple). (B) Frequency of mo-Macs and mo-DCs in the four conditions are depicted. Representative of 2 experiments with at least 3 biological replicates. Bars indicate mean ± SEM. Statistical significance was assessed by non-parametric Mann-Whitney test. *, P<0.05; **, P<0.01.

### NOD2 activates mTORC2 pathway and promotes anaerobic glycolysis

In an effort to further understand how NOD2 activation may impairs mTORC1-dependent macrophage differentiation, we then stimulated THP1 cells during 30 minutes with MDP and quantified the phosphorylation state of Regulatory-associated protein of mTOR (also known as Raptor) at serine 792 (S792) that is mediated by AMP-activated protein kinase (AMPK). This S792-phosphorylation of RAPTOR has been shown to reduce mTORC1 activity^41^ and to act as a metabolic checkpoint which coordinates the energy status of each cell^41^. In this experimental setting, activation of NOD2 signalling induced S792 phosphorylation of RAPTOR (SFig8A), suggesting that NOD2 could inhibit lipid uptake and foam cell formation through a negative feedback on mTORC1 activity. We next measured the phosphorylation of AKT at serine 473, as a surrogate marker of mTORC2 activation^40^. In agreement with our hypothesis, we observed that a short treatment with MDP of THP1 cells increased this specific phosphorylation of the residue S473 (SFig8B). This phosphorylation event is not observed in THP1 cells that lack the expression of NOD2 (SFig8B). Consistent with that observation, MDP-treated THP1 monocytic cells were characterized by a NOD2-dependent upregulation of IRF4 expression (SFig8C), which is a target gene of mTORC2^42^. At this time point, immunoblot analysis expectedly revealed that treatment with MDP did not induce the phosphorylation of S6K, which is a surrogate marker of mTORC1 activation (SFig8B). As mTORC2 pathway plays a key role in the glycolytic reprogramming of monocytes that are rapidly mobilized on demand^43,44^, we next investigated if MDP treatment of monocytes is associated with changes in mTORC2-mediated metabolic cascade. To this end, previously published RNAseq data for MDP-treated Ly6C^hi^ mouse monocytes (GEO accession number GSE101496) were mined for candidate genes encoding for enzymes involved in glycolysis. We observed a MDP-induced upregulation of several glycolytic genes such as the enzyme Gapdh that regulates the conversion of D-glyceraldehyde 3-phosphate into 1,3-bisphosphoglycerate (SFig9A)^20^. To get further insights how NOD2 may provide energy for monocytes, we next quantified the glycolytic capacity and reserve of THP1 cells that were deficient or not for NOD2. As THP1 cells mainly rely on glycolysis as a source of ATP for survival^45^, these cells represent a suitable model. Extracellular acidification rate (ECAR) was measured by using a Seahorse bioanalyser. During the first step of glycolysis, we noticed a similar basal glycolytic rate between NOD2-deficient and parental THP1 cells (SFig9B). Upon blockade of oxidative phosphorylation of ADP to ATP by oligomycin, the basal glycolytic capacity of wild-type cells was similar to the one of THP1 NOD2^-/-^ cells. By contrast, the inhibition of glycolytic H+ production by 2-Deoxy-D-glucose (2-DG) revealed a lower glycolytic reserve in the absence of NOD2. These results suggest that NOD2 may promote a lower pH, by sustaining a higher glycolytic demand, that is required for mo-DCs differentiation^14^. In agreement with the role of glucose on M-CSF-induced myelopoiesis^46^, we noticed a lowered expression of CD115 on THP1 cells that are deficient for NOD2 when compared to what is observed in controls (SFig9C,D). We then reasoned that NOD2 signalling may favour the conversion of Ly6C^hi^ to Ly6C^low^ monocytes, as observed for the addition of extracellular M-CSF^21^, by inhibiting mTOR pathway, which refrains the responsiveness to M-CSF of macrophage progenitor^21,46^. In line with the ability of NOD2 signalling to activate Signal transducer and activator of transcription 5 (STAT5)^47^, the treatment of THP1 cells with MDP significantly lowered the expression of IRF8 that is inhibited by STAT5 (SFig8C). Altogether, these results indicate that NOD2 signalling of monocytes may interfere with the PI3K/mTORC1 pathway through the mTORC2/AKT complex, which may inhibit diversion of monocyte differentiation to macrophages via metabolic reprogramming^43^. In other words, these data suggest that MDP induces a transient negative regulation of mTOR pathway to limit accumulation of inflammatory macrophages, leading to an unrepressed generation of mo-DCs.

### The inhibition of mTORC1 pathway by NOD2 promotes the secretion of TNF-alpha

We then hypothesized that activation of NOD2 signalling may influence the bioenergetic needs of nascent mo-DCs by interfering with the mTOR pathway. In agreement with the role of TNF-α on the development of mo-DCs, wortmannin increased by 3 fold the secretion of TNF-α upon stimulation of MDP-primed THP1 cells with LPS when compared to control cells (SFig10A, upper part). This effect of wortmannin on the monocyte responsiveness to LPS is lost with MDP-primed THP1 cells that do not express NOD2 (SFig10A, lower part). Conversely, the anti-inflammatory effect of wortmannin on the ability of terminally differentiated Nod2-deficient macrophages (PMA-Mac) to secrete TNF-α in response to LPS is lost when being treated concomitantly with MDP (SFig10C). Similarly to what was observed with wortmannin, rapamycin treatment did not impair the secretion of TNF-α by THP1 cells when stimulated with MDP (SFig10A). By contrast with what was observed with rapamycin, bafilomycin treatment, which inhibits autophagy at the final step of fusion of lysosome with autophagosome, blunted the synergistic effect of MDP (data not shown). We found that treatment with MHY1485 inhibited the MDP-induced secretion of TNF-α by THP1 cells (SFig10B). Altogether, these results suggest that Nod2 signalling on monocyte fate decision is dominant over the activation of mTORC1 for conditioning their differentiation into mo-DCs that are more responsive to bacterial stimulation as compared to macrophages. We next experimentally addressed the functional impact of early NOD2 signalling on the ability of nascent phagocytes to respond to LPS that is known to synergize with MDP^48^. As expected, these cells pre-treated with MDP enhanced their subsequent cytokine response upon LPS treatment (SFig10, left upper part). This synergistic effect on production of TNF-α to subsequent treatment with LPS was blunted with THP1 cells that do not express NOD2 (THP1 NOD2^-/-^) (SFig10A, lower part). In order to confirm our data with primary cells, mouse bone marrow monocytes were cultured with a growing media supplemented with MDP. The responsiveness to LPS was next analysed by measuring released TNF-α by specific ELISA. An apparent synergistic effect on secretion of TNF-α was retained on those cells that were primed with MDP (SFig11A). This synergistic effect on production of TNF-α to subsequent treatment with LPS was absent in monocytes that were isolated from the bone marrow of *Nod2*-deficient mice (SFig11B). Such a functional model of hierarchy could be of particular importance in a context of loss of bacterial tolerance or need for the replenishment of tissue mo-DCs following injury or infection.

### NOD2 loss-of-function mutation impairs the phenotypic switch of monocytes in CD patients in a TNF-alpha dependent manner

To evaluate the importance of a monocyte phenotypic switch in CD patients with NOD2 mutations, we use published RNA-seq dataset from an exploratory cohort that is deposited in the GEO database (GSE69446)^49^. Given that CD14-expressing cells are obligate precursors of discrete subsets of phagocytes that play a role in CD pathogenesis, the monocytes were isolated from peripheral blood of healthy controls (n=2) and CD patients in complete remission for at least 4 weeks prior to inclusion. Among those five patients, three carried the loss of function mutation in the *NOD2* gene, referred to as 1007fs mutation. As depicted in the Venn-diagram (Fig7A), MDP treatment of monocytes from healthy donors and CD patients significantly modified the expression of up to 362 and 1,660 genes respectively. Among those, a list of 306 genes were commonly regulated by MDP in blood monocytes from both control and CD patients. Geneset enrichment analysis (GSEA) indicated that the pathway “TNF-alpha signalling via NF-kB” (24/200)(adjusted p value 3.12E-20) was significantly induced among the 156 commonly differentially upregulated genes that are induced by MDP in CD14-expressing cells from either control or CD patients (Fig7B) (Supplementary data set 1). Among those, we noticed an enrichment of up-regulated genes related to dendritic cells (9/199)(adjusted p-value 7.8E-3) (Supplementary data set 2). Furthermore, a list of down-regulated genes that are related to diverse subsets of tissue macrophages was identified by using GSEA (11/204)(adjusted p-value 1.1E-5) (Supplementary data set 3). Among those patients, we next assessed whether cells bearing an unfunctional NOD2, as homozygous for the 1007fs NOD2 mutation may differentially respond to MDP. This led us to identify a specific loss of MDP-induced expression of 17 genes, including IL12B and miR-155 (Supplementary data set 4). Inhibition of miR-155 in human monocytes by an antagomir during 6h increased significantly MAFB^31^, a transcription factor implicated in the molecular control of monocyte-macrophage differentiation^50^. Pathway enrichment analysis identified a set of genes among the 17 that were implicated in TNF-alpha signalling via NF-kB (5/200)(adjusted p-value 2.53E-7)(Supplementary data set 5) and TNF-alpha effects on cytokine activity, cell motility, and apoptosis (4/135) (adjusted p-value 1.75E-4)(Supplementary data set 6). Accordingly, the Kyoto Encyclopedia of Genes and Genomes (KEGG) database analysis of the 941 up-regulated genes in CD cells which are not present in control cells after MDP treatment identified a set of genes related to “TNF signalling pathway” (28/112)(adjusted p-value 5.6E-11)(Supplementary data set 7). These results suggested that MDP inhibited the differentiation of monocytes into the macrophage pathway by the induction of DC-inducing soluble factors, such as TNF-α. We then evaluated whether the formation of mo-DCs that is initiated by NOD2 signalling is inhibited upon neutralization of TNF-α with adalimumab, which is a fully human anti-TNF-α monoclonal antibody. Interestingly, the percentage of classical monocytes was higher in patients responding to adalimumab than in patients not responding to the same drug^51^. CD14-expressing cells have been cultured for 5 days in a medium with GM-CSF, IL-4, MDP and/or adalimumab (Fig7C,D) and/or isotype control (data not shown). In contrast with the isotype control, the addition of adalimumab in the medium together with GM-CSF and IL-4 significantly increased the frequency of mo-Macs (Fig7D-E). Among those, it has been noticed that some co-express the macrophage marker CD14^52^. Interestingly, in the presence of MDP, mo-DCs did not express the marker CD14. Monocytes in the presence of MDP produced higher levels of TNF-α (SFig12). Morphological changes of the MDP-treated cells were observed by light microscopy as early as 24 hours after treatment with adalimumab. This effect was characterized by a lowered incidence of dendritic cell clustering (data not shown) and of mo-DCs as determined by flow cytometry (Fig7E). Interestingly, we observed that MDP is decreasing the level of CD115 as early as two days in adalimumab treated cells (Fig7C). In conclusion, NOD2 may molecularly define an education process that subsequently prevents accumulation of pathogenic macrophages.

**Figure 7.**
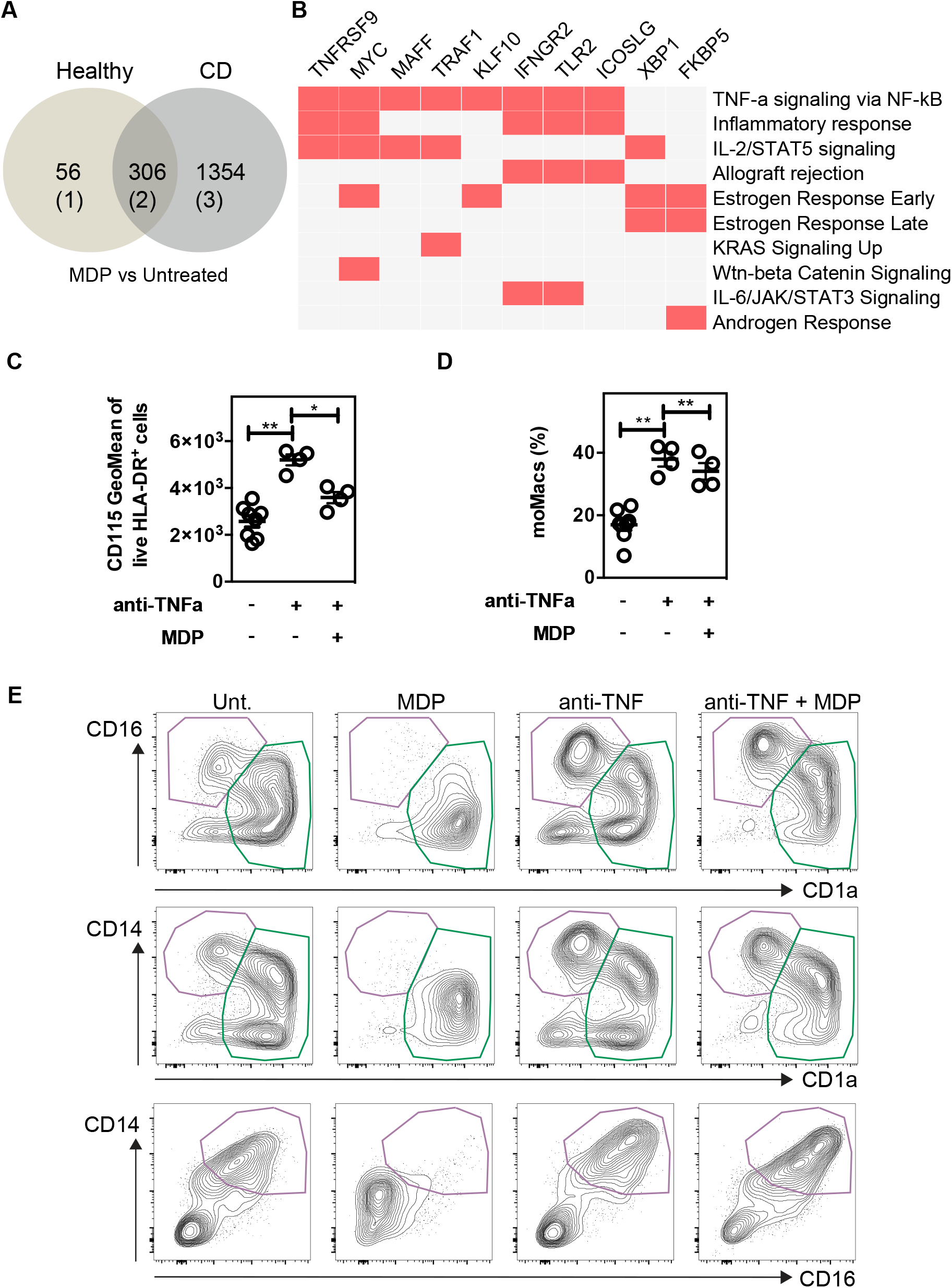
NOD2-induced decrease of macrophages is dependent on TNF-α production. Public RNA-seq dataset of untreated or MDP-stimulated monocytes isolated from peripheral blood of healthy controls (n=2) and CD patients in complete remission were analyzed. Among the CD patients, 3 are bearing and 2 are not bearing the loss of function mutation in the *NOD2* gene. (A) Venn diagram comparing the Differentially Expressed Genes (DEG) between MDP-treated and untreated in healthy controls and CD patients. (B) Geneset enrichment analysis (GSEA) among the 156 commonly differentially upregulated genes that are induced by MDP in CD14-expressing cells from either control or CD patients. (C) Monocytes were treated or not with MDP, anti-TNFα (Adalimumab) or both during GM/IL4 cultures. GeoMean of CD115 in live HLA-DR+ cells was measured at day 2 of culture. (D) Mo-DCs and mo-Macs frequencies were assessed by flow cytometry using the CD14, CD16 and CD1a markers at day 6 of the culture. The frequency of mo-Macs upon anti-TNFα and MDP treatment is depicted. (E) Contour representing the mo-DCs (green) and mo-Macs (purple). Representative of at least 2 independent experiments with different donors and with at least 4 biological replicates. Bars indicate mean ± SEM. Statistical significance was assessed by non-parametric Mann-Whitney test. *, P<0.05; **, P<0.01.

## Discussion

We report herein experimental evidence that monocytes fail to differentiate into mo-Macs when the Nod2-mediated signalling is activated. By using competitive bone marrow chimera, we did not observe a lack of recruitment of Ly6C^hi^ monocytes in the colon, suggesting that systemic MDP does not affect the number of recruited colonic monocytic cells, but may rather trigger early changes in epigenetic regulation of mo-DCs development^53^. Besides the Nod2-dependent regulation of GM-CSF secretion by stromal cells^23^, our data highlighted a Nod2-dependent regulation of a developmental process of mo-DCs by the gut microbiota that is likely solicited when the DCs population must be replenished. This is particularly true within the first years of life, in which the immunological tolerance is not yet fully operational. One may anticipate that such demand-driven generation of mo-DCs is likely dependent on several mechanisms governing tolerance to MDP that are programmed in time for keeping a fine balance between each discrete subsets of phagocytes with context-dependent functions. Chronic stimulation of monocytes with MDP causes what is often called NOD2-induced tolerance that consists in a tolerance to MDP and other bacterial signals such as LPS^54^. Proteasomal degradation of NOD2 protein confers rapid induction of refractoriness to MDP that protects the host from tissue damage or even death^54, 55^. Such negative feedback regulatory mechanism fails to occur when monocytes are defective in either the E3 Ubiquitin ligase ZNRF4 or the protein NLRP12^56, 57^. Consequently, treatment with MDP of monocyte-derived cells that are deficient for the aforementioned molecules led to an excessive inflammation with a sustained NF-κB activation. Different studies have shown that most of the immune cells, particularly myeloid cells, may actually have a dual activity, pro-inflammatory or immunosuppressive, depending on the signals received from the tumour microenvironment^58^. As this dual display of opposing antagonising functions exists also in conventional dendritic cells^59^, we anticipated that similar dual properties of mo-DCs would likely exist and be influenced by microbial-derived products in their microenvironment.

The notion that metabolic control is upstream of inflammatory function has been proposed recently^60^. Indeed, circulating monocytes and monocyte-derived macrophages from patients with fibroinflammatory vasculopathy are highly efficient in glucose import and are expressing higher glycolysis-associated genes (GLUT1, HK2, PKM2, LDH, c-myc, and HIF-1α) in comparison to healthy individuals. Consistently, the expression of the Aldoa and Aldoc that are converting F1,6BP into GADP and of the Gadph that is converting GADP into 1,3BPG were significantly upregulated in monocytes that were treated by MDP (SFig9A)^20^. Similarly, MDP stimulation enhanced the expression of the Pgk1 converting 1,3BPG into 3-P-G and the Pagm1 that is involved in the conversion of the latter into 2PG. Aside from these enzymes, the Ldhb which is converting the Lactate into Pyruvate was downregulated, as well as the HK-II which is converting the glucose into glucose 6-phosphate (G6P). Among the 156 up-regulated genes induced by MDP in control and CD cells, analysis of MSigDB database indicated also the pathway “mTORC1 signalling” and containing SLC7A5, XBP1, HSPA5, PPA1, AK4 and CFP. Further studies will be needed to better understand the MDP-induced regulation of glycolysis enzymes and their regulation of mTORC1 signalling^61^. Additionally, among the genes having a loss of MDP-induced expression in monocytes from SNP13 CD patients, *AK4* is involved in the positive regulation of mouse myeloid cells glycolysis and inflammatory cytokine production such as Tnf-α and Il-6^62^. As microbiota-derived circulating peptidoglycan is found in the mouse blood^63^, one can propose that MDP can induce a metabolic reprogramming of circulating monocytes leading to a higher glucose consumption which regulates their mitochondrial activity, such as ROS production and effector molecules expression such as TNF-α.

Given that MDP treatment lowers the expression level of CD115 at an early stage of the differentiation process of monocytes into mo-DCs, we presumed that this phenomenon might rely on the responsiveness of mo-DCs progenitors to TNF-α. Although it is not known whether those balancing feedbacks may foster the development of mo-Macs, we demonstrated here that blocking TNF-α with adalimumab arrests the MDP-induced development of mo-DCs. This is in accordance with the upregulation of M-CSFR expression observed in response to adalimumab, which has been described to bind membranous TNF-α with a relatively higher affinity than etanercept^64^. Interestingly, DC obtained from either *Tnfr1*^-/-^ mice or patients treated with anti-TNF-α showed an unusual mixed immature/mature phenotype^65, 66^, suggesting the development of macrophages in these cultures^27, 67^. While TNF-α is a weak stimulator of CCR7 expression^68^, it counterbalances the emergence of M2-like tumour macrophages^69^. Such cytokine is sharply induced by microbiota, in Nod2-dependent and –independent pathways, during weaning for lowering risk of developing colorectal cancer later in life^64^. Conversely, an impaired dendritic cell function has been reported in most CD patients with NOD2 1007fs mutation^70^. While TNF-α can upregulate NOD2 expression in myelomonocytic cells^39^, anti-TNF therapy could alter the Nod2-induced equilibrium between discrete subsets of intestinal phagocytes with different properties. Additional analysis of published RNAseq data indicated that the pathway “Wnt-beta Catenin Signalling” was also significantly induced among the 156 common genes induced by MDP in control and CD cells including MYC, HEY1, JAG1 and A Disintegrin And Metalloproteinase 17 (ADAM17). Wnt-beta catenin signalling pathway limits the differentiation into macrophages of bone marrow cells cultured with GM-CSF^42^. It is worth noting that A Disintegrin And Metalloproteinase 17 (ADAM17 (also known as TNF-alpha converting enzyme (TACE)) is a sheddase with a broad range of substrates such as membrane bound TNF-α^71^ and M-CSF receptor^32^. ADAM17-dependent cleavage of M-CSF receptor is the mechanism by which GM-CSF and IL-4 block M-CSF- and RANKL-induced osteoclast differentiation from monocytes^32^. Modulating the NOD2/TNF-α signalling axis to balance induction of mo-DCs and repression of mo-Macs appears to be a promising new target for immunotherapy of colorectal cancer and to treat stricturing complications of CD patients.

Additionally, among the 17 genes losing MDP-induced expression in monocytes from CD patients with the homozygous SNP13 mutation, *NR4A3* is involved in the proper differentiation of Mo-DCs (FC 1.72, adjusted p-value 0.00206)^72^. One may suggest that inhibition of M-CSF receptor signalling by MDP on monocytes is required to impaired macrophage differentiation in certain circumstances. For instance, type 1 cysteinyl leukotriene receptor (CYSLTR1) was significantly down-regulated from healthy controls or CD patients under MDP treatment^49^. Inhibition of CYSLTR1 prevents M-CSF- and RANKL-induced osteoclast differentiation of bone marrow precursors^73^. Additionally, the ETS variant transcription factor 3 (ETV3) was significantly up-regulated within the MDP-treated monocytes from healthy controls or CD patients. It is induced by the anti-inflammatory cytokine IL-10^74^, and blocks M-CSF-induced macrophage proliferation^75^. In CD patients, a unique response to MDP was observed with 941 and 413 up-regulated and down-regulated genes respectively^49^. Analysis of Azimuth Cell Type database showed an enrichment of up-regulated genes that are related to “Myeloid Dendritic Type 1” among which CCR7 is a hallmark of DC as it is critically required for their migration to lymph nodes^76^. Equally of importance, the transcription factor *NRA43*, that is involved in the proper differentiation of Mo-DCs^72^, is less induced by MDP treatment in monocytes from CD patients than in cells from healthy controls (FC 1.6 and FC 1.71, respectively) and even less in cells from patients with loss-of-function alleles of *NOD2* in comparison to control cells under the same treatment. Here, we provided a better understanding of macrophage-differentiation inhibition by NOD2-mediated signalling, which may offer new therapeutic strategies aiming at limiting detrimental effects of pathogenic macrophages in gut pathologies such as CD.

## Supporting information

Supplementary data set 1

Supplementary data set 2

Supplementary data set 3

Supplementary data set 4

Supplementary data set 5

Supplementary data set 6

## Author Contributions

Conceptualization: CC, DA, ES, MC and LFP. Methodology: CC, DA, JK, KR, OB, NW, MD, ES, MC and LFP. Formal analysis: JK, WL, NW, KR, MC, LFP. Investigation: CC, DA, KR, OB, NW, MD, MC and LFP. Writing – original draft: MC and LFP. Writing – review and editing: all authors. Visualization: CC, DA, MC and LFP. Supervision: LFP and MC. Funding acquisition: MC and LFP.

## Acknowledgements

This work was funded by the French government’s ATIP-Avenir program. LFP also received a fellowship from the ATIP-Avenir program and funding support by the French national IBD patients’ association (Association François Aupetit (AFA). CC received a fellowship funded by the cancer charity “La Ligue contre le cancer”. ES received a funding support by the Agence Nationale de la Recherche (grant number ANR-17-CE15-0011-01). We thank the staff at the animal and cytometry facility at the Pasteur Institute of Lille. Pathway analysis has been realised with Enrichr^77-79^.

## Competing interests

The authors declare no competing interests.

## Materials and Methods

### Mice

All animal studies were approved by the local investigational review board of the Institut Pasteur of Lille (N°28010-2016012820187595). Animal experiments were performed in an accredited establishment (N° B59-108) according to governmental guidelines N°86/609/CEE. Age-matched and gender-matched C57BL/6J WT, Nod2-deficient mice (Nod2^-/-^), Nod2ΔLyzM, Nod2^fl/fl^ have free access to standard laboratory chow diet in a temperature-controlled SPF environment and a half-daylight cycle exposure. C57BL/6J WT GF, Nod2^-/-^ GF mice were bred at TAAM-CNRS and were transferred into autoclaved sterile micro-isolator cages. C57BL/6J WT mice were purchased from Janvier Laboratories, France. *Nod2*^*–/–*^ mice were provided by R.A. Flavell (Yale University School of Medicine, Howard Hughes Medical Institute). We thank Dr Philip Rosenstiel (Institute of Clinical Molecular Biology, Kiel, Germany) for providing the Nod2^fl/fl^ mice, and the Jackson (stock #004781) for the LyzM-Cre mice^34^. Conditional knockout of Nod2 in the lysozyme M expressing cells was established by crossing *LyzM-*Cre with *Nod2*^fl/fl^ mice, resulting in *Nod2*^ΔLyzM^ or *Nod2*^fl^.

### Induction of Relapsing-Remitting Colitis

Relapsing-remitting colitis was induced by giving mice 2% (wt/vol) DSS (TdB Consultancy) for a period of 5 days followed by normal drinking water for 7 days with a threshold of maximal lost weight of 20% of the initial weight. DSS was dissolved in drinking water and changed every 3 days. Signs of morbidity, including body weight, stool consistency, and occult blood or the presence of macroscopic rectal bleeding, were checked daily. At specific time points throughout the course of the challenge, mice were autopsied to assess the severity of the disease by measurement of colon lengths and cell composition by flow cytometry. In some conditions, mice were treated with 200 ppm indole-3-carbinol (Sigma) during 4 weeks before the cycle of DSS as described previously^8^. In some experiments, mice were injected with either Nod2^+/+^ or Nod2^-/-^ Ly6C^hi^ monocytes (1×10^6^ cells/mice) 4.5 days after induction of the colitis protocol.

### Bone marrow transplantation experiments

Recipient mice underwent a lethal total-body irradiation (2X 5.5Gy, 4h between each dose). Twenty-four hours post-irradiation, mice received intravenously 2 × 10^6^ fresh bone marrow cells. *Nod2*-deficient animals were irradiated and reconstituted in a 1:1 ratio with bone marrow cells from WT (CD45.1) and *Nod2*-deficient mice (CD45.2). Blood was collected in heparin-containing tubes 7–8 weeks after bone-marrow transplantation and reconstitution efficiency was checked by flow cytometry^80^. Cellular content within the colon, the blood and the peritoneum of chimeric mice was analyzed 8 weeks after bone marrow reconstitution by flow cytometry. In some experiments, Dextran sodium sulfate (DSS) was administered for 5 days in the drinking water of mixed-bone marrow chimera mice to induce acute colitis.

### Fecal transplantation

Fecal microbiota from wild-type mice was transplanted by gavage in germ-free mice that are deficient or not for Nod2^81^. The mice were used after four weeks of colonization with feces from wild-type mice.

### Isolation of mouse colonic lamina propria cells

Lamina propria Mononuclear Cells (LPMC) were prepared from murine intestines by enzymatic digestion as previously described^3^. Briefly, cells were isolated from colons, after removal of epithelial cells, by enzymatic digestion with 200mg/ml fungizone, 1.25 mg/ml collagenase D (Roche Diagnostics), 0.85 mg/ml collagenase V (Sigma-Aldrich), 1 mg/ml dispase (Life Technologies), and 30 U/ml DNaseI (Roche Diagnostics) in complete RPMI 1640 for 30–40 min in a shaking incubator until complete digestion of the tissue. After isolation, cells were passed through a 40μm cell strainer before use (BD biosciences). Colonic cell numbers were determined with counting beads and following manufacturer’s instructions (AccuCheck counting beads, Invitrogen).

### Generation of bone marrow-derived dendritic cells and macrophages

Bone marrow cells were flushed out of the mouse bones with complete RPMI 1640 (Gibco). A single-cell suspension was then prepared by repeated pipetting. Bone marrow-derived macrophages (BMDMs) and dendritic cells (BMDCs) were generated for 7 days in respectively IMDM or RPMI-1640 medium (Gibco), supplemented with glutamine, penicillin, streptomycin, 2-mercaptoethanol ([all from Gibco)], and 10% heat-inactivated fetal calf serum (GE Healthcare). The medium was supplemented either with 20% supernatant of L929 cells (M-CSF-producing cells) for BMDMs or with 20 ng/ml GM-CSF of J558 cells (GM-CSF-producing cells) for BMDCs. For the latter, half of the medium was removed at day 3 and new medium supplemented with GM-CSF supernatant (2x, 40 ng/ml) was added^27^.

### Human monocyte-derived dendritic cell generation

Human PBMC were prepared from buffy coats (Etablissement Francais du Sang (EFS), Lille, France) using Ficoll Paque (Lymphoprep, StemCell). The use of human samples was approved by the French Ministry of Education and Research under the agreement DC 2013-2575. According to French Public Health Law (art L 1121–1-1, art L 1121– 1-2), Institutional Review Board and written consent approval are not required for human non-interventional studies. Monocytes were positively isolated using CD14+ microbeads (Miltenyi Biotec) according to the manufacturer’s recommendations. Cells were cultured for 6 days in GM-CSF (20ng/ml; Peprotech) and IL-4 (5ng/ml; Peprotech). When mentioned, Muramyl dipeptide (MDP) (10μg/ml; Invitrogen) was added from the beginning of the culture. The mTOR activator MYH1485 (0.5μM, Sigma) was added at the start of the culture. Adalimumab (Humira M02-497) was a gift from Abbott (Abbott Park, IL, USA).

### Cytokine measurement

Cytokine levels were determined by ELISA kits (DuoSet), according to protocols provided by R&D Systems.

### Western blot

Protein extraction was performed using RIPA buffer in the presence of complete Mini EDTA-free protease inhibitor (Roche) and PhosSTOPTM phosphatase inhibitor (Roche). Protein separation was performed by SDS-page using Bolt 4 to 12% Bis-Tris protein gels (Invitrogen). Transferences were done in an iBlot 2 gel transfer device using iBlot 2 transfer nitrocellulose stacks (Invitrogen). Membranes were blotted against phospho-AKT (Ser473) (Cell Signaling), phospho-RAPTOR (Ser792) (Cell Signaling), phospho-p70 S6 Kinase (Thr389) (Cell SSignaling), β-ACTIN (Cell Signaling) and their correspondent HRP-conjugated secondary antibodies. The revelation was performed using the SuperSignal West Femto Maximum Sensitivity Substrate (Thermo Scientific) and images were acquired using an ImageQuant LAS 4000 (GE Healthcare).

### Flow cytometry

Single-cell suspensions were stained and analysed using a FACS LSRFortessa™ system (BD Biosciences). Dead cells were excluded with the LIVE/DEAD Fixable Violet Dead Cell staining kit (Life technologies). The cells were then incubated for 10 minutes with purified rat anti-mouse CD16/CD32 (Biolegend, 93 clone)(only for mouse cells) and normal mouse serum (Interchim) before being stained with various monoclonal antibodies for 20 minutes in the dark on ice. For mouse cells, lineage-positive cells were excluded using the PerCP5.5-conjugated anti-CD3 (17A2), anti-NK1.1 (PK136), anti-CD19 (6D5), anti-Ly6G (1A8) (Biolegend). PerCP-conjugated anti-CCR3 (83103) added to the lineage staining to exclude eosinophils was from R&D. Alexa Fluor 700-conjugated anti-Ly6C (AL21) was from BD Pharmingen. PECF594-conjugated anti-CD11c (HL3) was from BD Horizon. Allophycocyanin-Cy7-conjugated anti-CD11b (M1/70), Brilliant violet 510-conjugated anti-MHC Class II (I-A/I-E) (M5/114.15.2), Allophycocyanin-conjugated anti-CD64 (X54-5/7.1), PE-Cy7-conjugated anti-CD24 (M1/69), Brilliant violet 650-conjugated anti-CD45.2 (104), Brilliant violet 711-conjugated anti-CD45.1 (A20), PE-conjugated anti-CD226 (10E5), APC-conjugated anti-CD115 (AFS98) and FITC-conjugated anti-CD102 (3C4) were all from Biolegend. PE-conjugated anti-CCR2 (475301) was from R&D systems. The data were analysed with Flowjo software V10.1 (TreeStar). For human cells, similar procedure was used with anti-HLA-DR FITC (eBioscience, clone LN3), anti-CD1a APC (Biolegend, clone HI149), anti-CD16 PE-Cy7 (BD Pharmingen, clone 3G8), anti-CD14 PE (Miltenyi Biotec, clone REA599). Surface CD115 (M-CSFR) was detected using rat anti-CD115 (Santa Cruz, clone 3-4A4) revealed by a Alexa Fluor 488-labelled rabbit anti-rat IgG (Thermofisher)^52^

### THP-1 cell culture and stimulation

The THP-1 monocytic cell line was cultured in RPMI 1640 medium (Gibco) supplemented with 10% heat inactivated FBS (Gibco), L-glutamine (Thermo Fisher), MEM non-essential amino acids (Thermo Fisher), sodium pyruvate (Thermo Fisher), HEPES (Thermo Fisher) and 0.05mM of 2-mercaptoethanol (Thermo Fisher). Cells were kept in culture at cell concentrations ranging from 2×10^5^ cells/mL to 8×10^5^ cells/mL and routinely verified negative for mycoplasma contamination by PCR analysis. THP-1 stimulation was performed in 96-well flat bottom plates at 1×10^5^ cells per well in a final volume of 200 μl. Cells were stimulated with two sequential treatments of 24 hours each. For the first 24 hours of treatment, cells were cultured in RPMI complete medium or RPMI medium with MDP at 100μg/ml. The second 24 hours treatment consisted of LPS at 50ng/ml or RPMI medium. In selected experiments the first treatment was MHY1485 (Sigma) at 2μM, rapamycin (Sigma) at 100, or wortmannin (Sigma) at 2 μM. In some conditions, THP-1 macrophages were generated by adding PMA (5ng/ml) for 48h, followed by at least 2 days without PMA^35,82^. Culture supernatants were collected after the second treatment and TNF-α levels were quantified by ELISA using the Human TNF-alpha DuoSet ELISA (R&D systems) following manufacturer recommendations.

### Extracellular acidification rate (ECAR)

ECAR was measured under basal conditions and in response to glucose (10mM) using the Seahorse Glycolysis Stress Test Kit by using a Seahorse bioanalyser.

### Gene expression

RNAs were extracted using the RNEasy mini kit (Qiagen). Isolated RNA was reverse-transcribed with the cDNA synthesis kit (Agilent Technologies), according to the manufacturer’s instructions. The resulting cDNA (equivalent to 500ng of total RNA) was amplified using the SYBR Green real-time PCR kit and detected on a Stratagene Mx3005 P (Agilent Technologies). qPCR was conducted using forward and reverse primers (sequences available upon request). Relative abundance of gene expression was assessed using the 2-ΔΔCt method. Actb was used as an internal reference gene in order to normalize the transcript levels.

### Statistics

Data were analyzed using Prism6.0 (GraphPad Software, San Diego, CA). Statistical significance was assessed by non-parametric Mann-Whitney test or two-way ANOVA for multiple comparisons. Values represent the mean of normalized data ± SEM. *, P<0.05; **, P<0.01; ***, P<0.001; ****, P<0.0001.

## Notes

### Competing Interest Statement

The authors have declared no competing interest.

