## Supplementary Figures for "Monocyte intrinsic NOD2 signalling inhibits pathogenic macrophage differentiation and its loss in inflammatory macrophages improves intestinal inflammation"

**Supplementary figure legends**

**Figure supp1.** Fecal microbiota from wild-type mice was transplanted in germ-free mice that are deficient or not for NOD2. Four weeks after colonization, the proportions of mononuclear phagocytes were evaluated in the transplanted mice. (A) Experimental set-up. (B) Frequency and absolute number of CD11c- MHCII+ and CD11c+ MHCII+ cells (3 mice per group).

**Figure supp2 related to figure 1.**Gating strategy in WT GF (A), in Nod2-/- SPF, and in Nod2-/- GF as explained in Figure 1. Contour plots and the frequency of conventional DC1, DC2, of mo-DCs, mo-Macs and Monocyte MHCII+ are depicted.

**Figure supp3 related to figure 2.** Mixed bone marrow chimera mice were generated as described in Figure 2. Absolute numbers (A) and frequency (B) of total WT and Nod2-/- mo-DCs and mo-Macs in the colon of recipients are depicted and the ratio of mo-DCs vs mo-Macs (4 mice). Bars indicate mean ± SEM. Statistical significance was assessed by non-parametric Mann-Whitney test. * $P<0.05$.

**Figure supp4 related to figure 2.** Gating strategy of the mixed bone marrow chimera mice generated as described in Figure 2G. (A) Contour plots of conventional DC1, DC2, mo-DCs, CD11c+ Macs, mo-Macs, Monocyte MHCII+ and Monocyte MHCII- are depicted. (B) Each subset is represented according to CD45.1 and CD45.2. (C) MHCII GeoMean in Monocyte MHCII+.

**Figure supp5.** Expression of NOD2 in mo-DCs upon differentiation for 5 days with GM-CSF and IL-4. Bars indicate mean ± SEM from three biological replicates. Statistical significance was assessed by multi-comparison non-parametric Friedman paired test, with Dunn's post test. $P < 0.05$ (*), $P < 0.01$ (**).

**Figure supp6 related to figure 5.** Relative expression of *Nod2* in splenic CD4+ T cells, peritoneal macrophages (Mac), and M-CSF generated bone-marrow macrophages (BMDM) in Nod2ΔLyzM (Lyz2Cre/+) and in littermate control flox animals (Lyz2+/+). Bars indicate mean ± SEM at least three mice per group**.**

**Figure supp7.** Wild-type (blue) and Nod2-/- (red) mice were fed with a diet enriched for indole-3-carbinole (I3C), an AhR agonist (open circle), or with control diet (closed circle) during 4 weeks before induction of DSS-mediated colitis. Body weight loss was measured over a 6-day course of 2% DSS. Data are representative of five to six mice per group. Bars indicate mean ± SEM. Statistical significance was assessed by two-way ANOVA, Bonferroni's multiple comparisons test. **, $P<0.01$; ***, $P<0.005$, ****, $P<0.001$.

**Figure supp8.** (A) THP1 (left part) and THP1 *NOD2*-deficient cells (right part) were stimulated for the indicated time with MDP or MDP then LPS and the presence of phosphorylated RAPTOR (Ser792) was measured by western blot, and the loading controlled by β -ACTIN measurement. (B) THP1 (left part) and THP1 *NOD2*-deficient cells (right part) were stimulated for the indicated time with MDP, LPS or both and the presence of phosphorylated AKT (Ser473) and phospho-p70 S6 Kinase (Thr389) was measured by western blot, and the loading controlled by β-ACTIN measurement. (C) *IRF4* and *IRF8* mRNA expression in THP-1 (blue) and THP-1 NOD2-/- (red) monocytic cell line was measured by RT-qPCR at the beginning of the culture or 24h and 48h after MDP treatment. Data are representative of 2 independent experiments with at least three biological replicates. Bars indicate mean ± SEM. Statistical significance was assessed by ordinary one- way multiple comparisons (A). ***, P<0.005, ****, P<0.001.

**Figure supp9.** MDP enhances the differentiation of Mo-DCs in a glycolytic and MTORC1 independent manner. (A) Overexpressed enzymes involved in glycolysis in MDP-treated mouse monocytes in published RNA-seq datasets (GEO accession number GSE101496). (B) Extracellular acidification rate (ECAR) was measured in the MDP-treated Nod2+/+ (blue) and Nod2-/- THP-1 (red) cells. Data are representative of 2 independent experiments with at least four biological replicates. (C-D) Expression of CD115 (GeoMean) on Nod2+/+ (blue) and Nod2-/- THP-1 (red) cells was measured by flow cytometry (C) and visualised by confocal analysis (ImageStream) (D). Bars indicate mean ± SEM. Statistical significance was assessed by non-parametric Mann-Whitney test. * P<0.01.

**Figure supp10.** (A) THP1 (blue) and THP1 *NOD2*-deficient cells (red) were treated as described in material and method section to evaluate LPS responsiveness of MDP-treated cells. In addition, the mTORC1 activator wortmannin or the mTOR inhibitor rapamycin were added or not for 24h. TNF-α production was measured. (B) LPS responsiveness was evaluation by measuring TNF-α production in THP1 WT or Nod2-/- cells treated with MDP, MHY1485 or both for 24h. (C) Differentiated macrophages with PMA (PMA-Mac), expressing or not NOD2, were treated as described above with wortmannin and TNF-α production was measured. (D) MDP responsiveness of LPS-treated PMA-Mac was evaluated as described above. Bars indicate mean ± SEM at least three biological replicates and data are representative of 2 independent experiments. Statistical significance was assessed by ordinary one-way multiple comparisons. *, P<0.05., **, P<0.01, ***, P<0.005 ****, P<0.001.

**Figure supp11.** Mouse bone marrow monocytes from WT (A) and Nod2-/- mice (B) were treated with MDP or LPS for 24h ($1^{st}$ Tx), and washed before a second treatment with MDP or LPS (2d Tx). mTNF-α was measured 24h after the last treatment in the supernatant. Data are representative of 2 independent experiments with at least three biological replicates. Bars indicate mean ± SEM. Statistical significance was assessed by non-parametric Mann-Whitney test. * P<0.01.

**Figure supp12.** Mo-DCs were differentiated in the presence of MDP, and hTNF-α was measured at 24h. Bars indicate mean ± SEM of four biological replicates. Statistical significance was assessed by non-parametric Mann-Whitney test. * P<0.05.

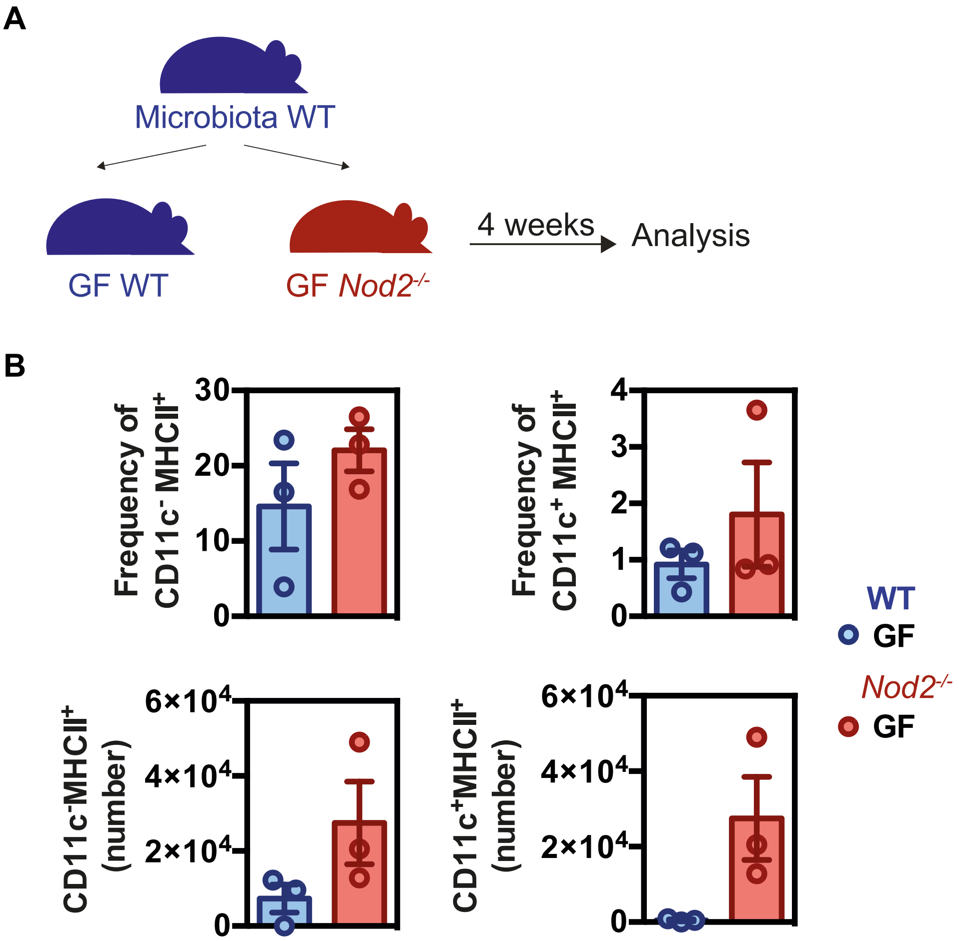
A
Microbiota WT
GF WT
GF Nod2-/-
4 weeks
Analysis
B
Frequency of CD11c- MHCII+
Frequency of CD11c+ MHCII+
CD11c-MHCII+ (number)
CD11c+MHCII+ (number)
WT
GF
Nod2-/-
GF

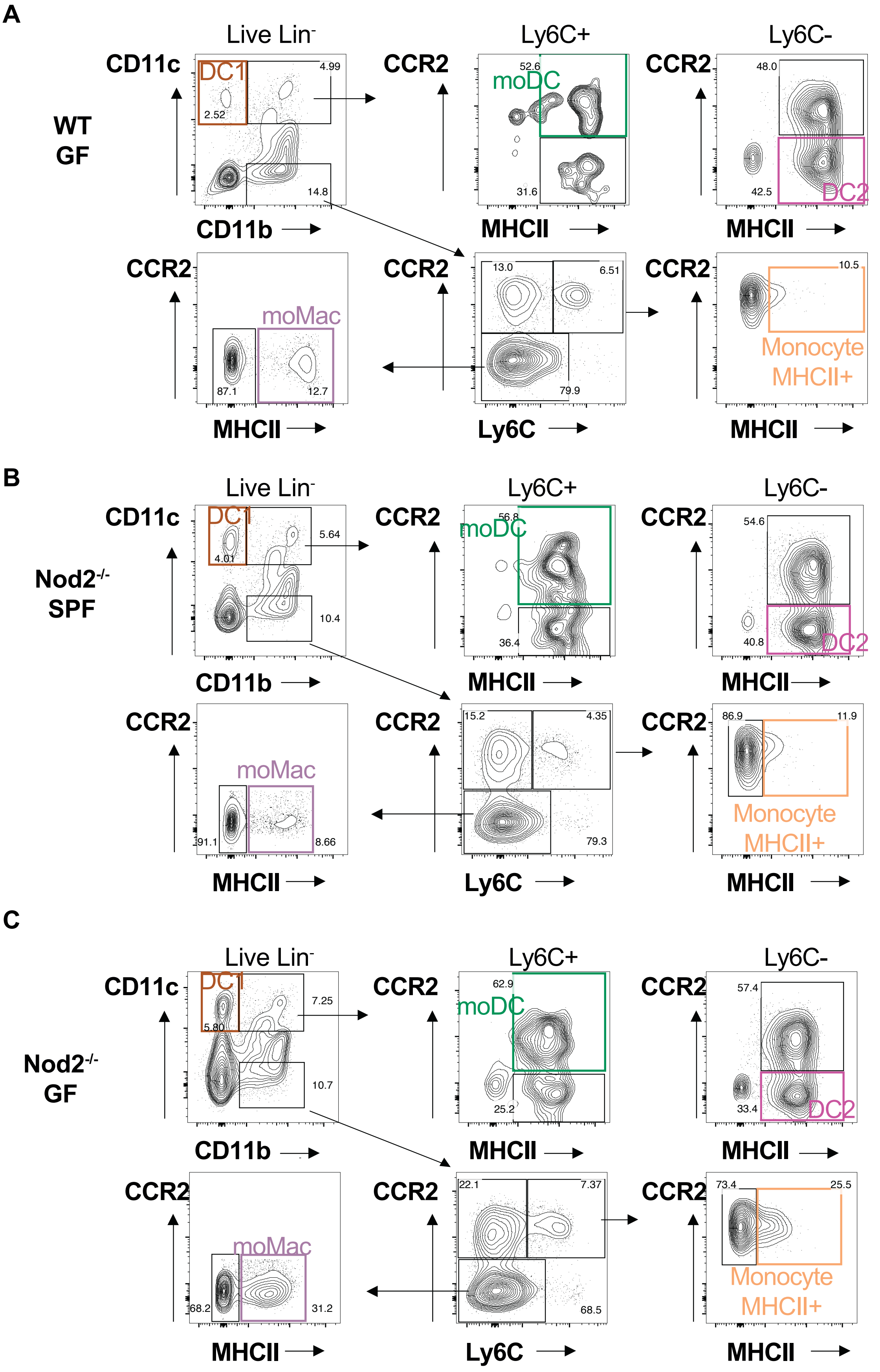
A
WT GF
Live Lin-
CD11c
DC1
2.52
4.99
14.8
CD11b
Ly6C+
CCR2
52.6
moDC
31.6
MHCII
Ly6C-
48.0
42.5
DC2
CCR2
moMac
87.1
12.7
13.0
6.51
79.9
Ly6C
10.5
Monocyte MHCII+
B
Nod2-/- SPF
4.01
5.64
10.4
56.8
36.4
54.6
40.8
91.1
8.66
15.2
4.35
79.3
86.9
11.9
C
Nod2-/- GF
5.80
7.25
10.7
62.9
25.2
57.4
33.4
68.2
31.2
22.1
7.37
68.5
73.4
25.5

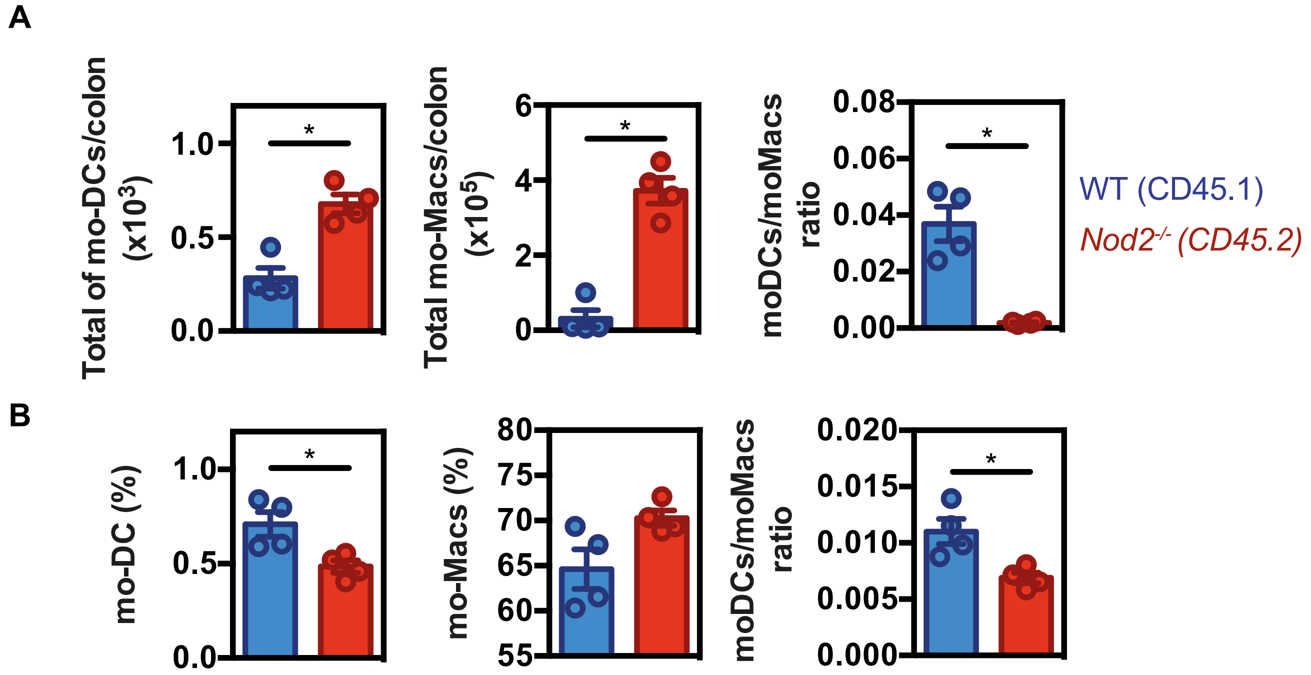
A
Total of mo-DCs/colon (x10^3)
1.0
0.5
0.0
*
Total mo-Macs/colon (x10^5)
6
4
2
0
*
moDCs/moMacs ratio
0.08
0.06
0.04
0.02
0.00
*
WT (CD45.1)
Nod2-/- (CD45.2)
B
mo-DC (%)
1.0
0.5
0.0
*
mo-Macs (%)
80
75
70
65
60
55
moDCs/moMacs ratio
0.020
0.015
0.010
0.005
0.000
*

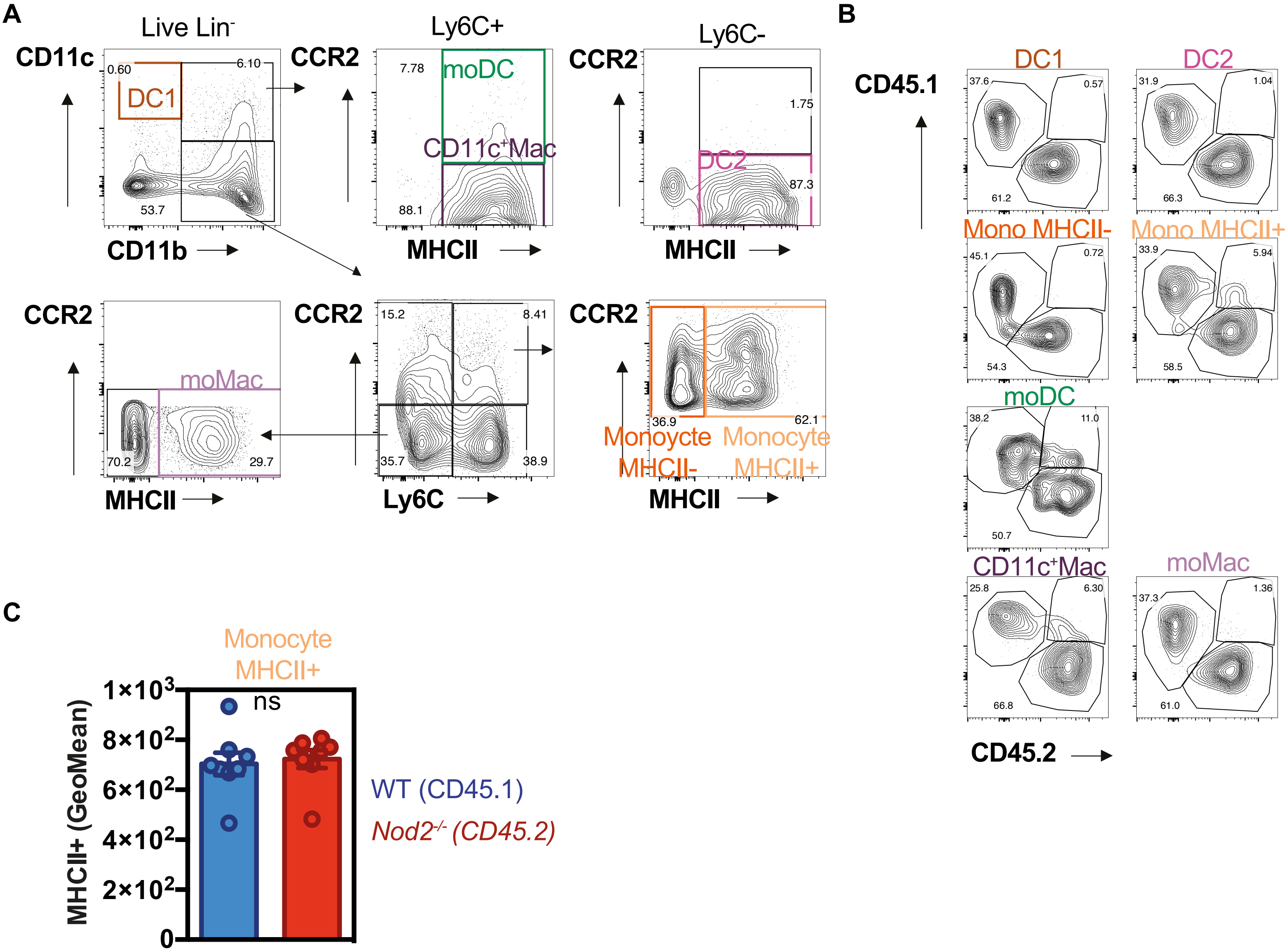
A
Live Lin-
CD11c
0.60
DC1
6.10
53.7
CD11b
Ly6C+
CCR2
7.78
moDC
CD11c+Mac
88.1
MHCII
Ly6C-
CCR2
1.75
DC2
87.3
MHCII
CCR2
moMac
70.2
29.7
MHCII
CCR2
15.2
8.41
35.7
38.9
Ly6C
CCR2
36.9
62.1
Monoycte MHCII-
Monocyte MHCII+
MHCII
B
CD45.1
DC1
37.6
0.57
61.2
DC2
31.9
1.04
66.3
Mono MHCII-
45.1
0.72
54.3
Mono MHCII+
33.9
5.94
58.5
moDC
38.2
11.0
50.7
CD11c+Mac
25.8
6.30
66.8
moMac
37.3
1.36
61.0
CD45.2
C
Monocyte MHCII+
MHCII+ (GeoMean)
1×10³
8×10²
6×10²
4×10²
2×10²
0
ns
WT (CD45.1)
Nod2-/- (CD45.2)

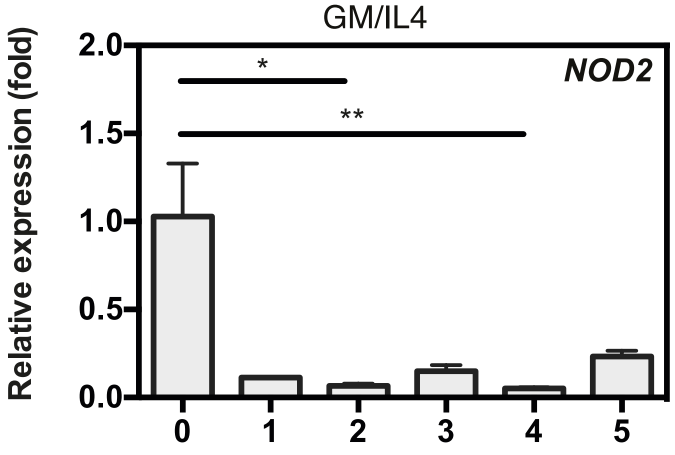
GM/IL4
NOD2
Relative expression (fold)
2.0
1.5
1.0
0.5
0.0
*
**
0
1
2
3
4
5

Supplementary Figure 6

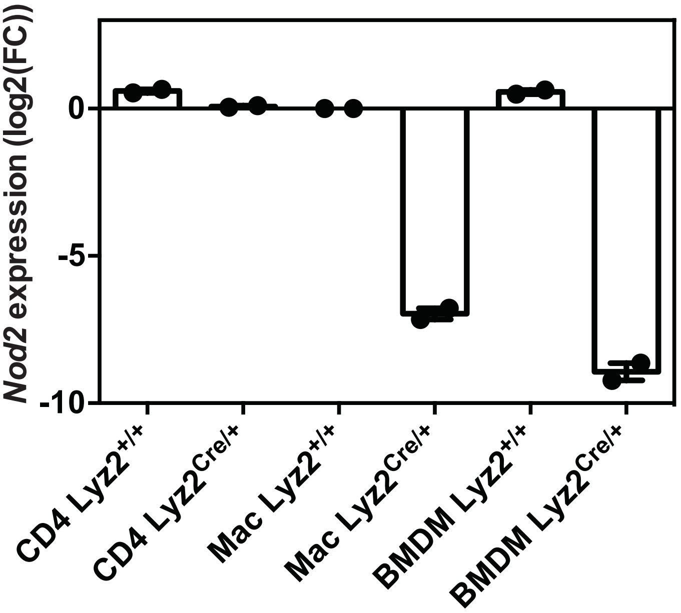

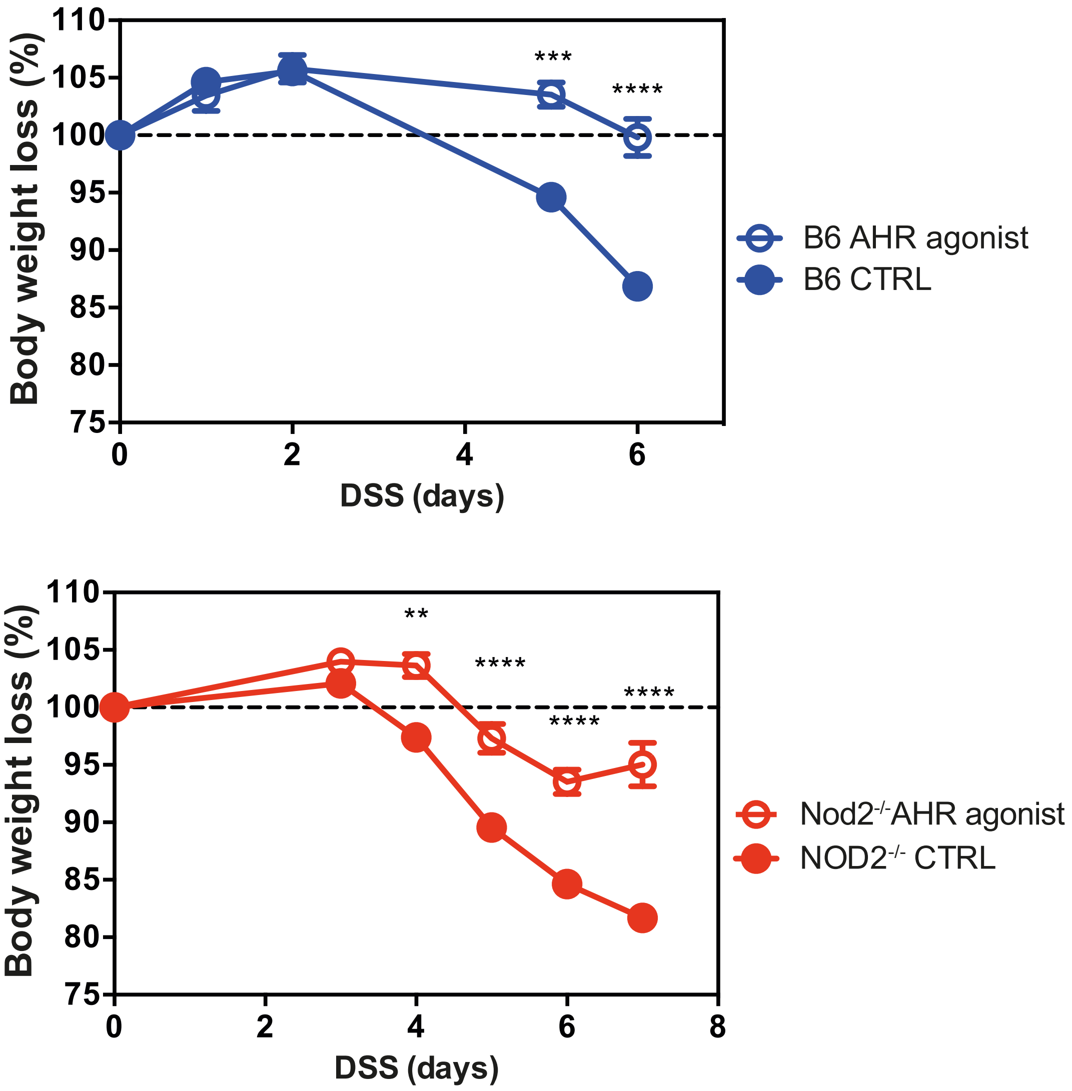

Body weight loss (%)
110
105
100
95
90
85
80
75
0
2
4
6
DSS (days)
***
****
B6 AHR agonist
B6 CTRL
Body weight loss (%)
110
105
100
95
90
85
80
75
0
2
4
6
8
DSS (days)
**
****
****
****
Nod2-/-AHR agonist
NOD2-/- CTRL

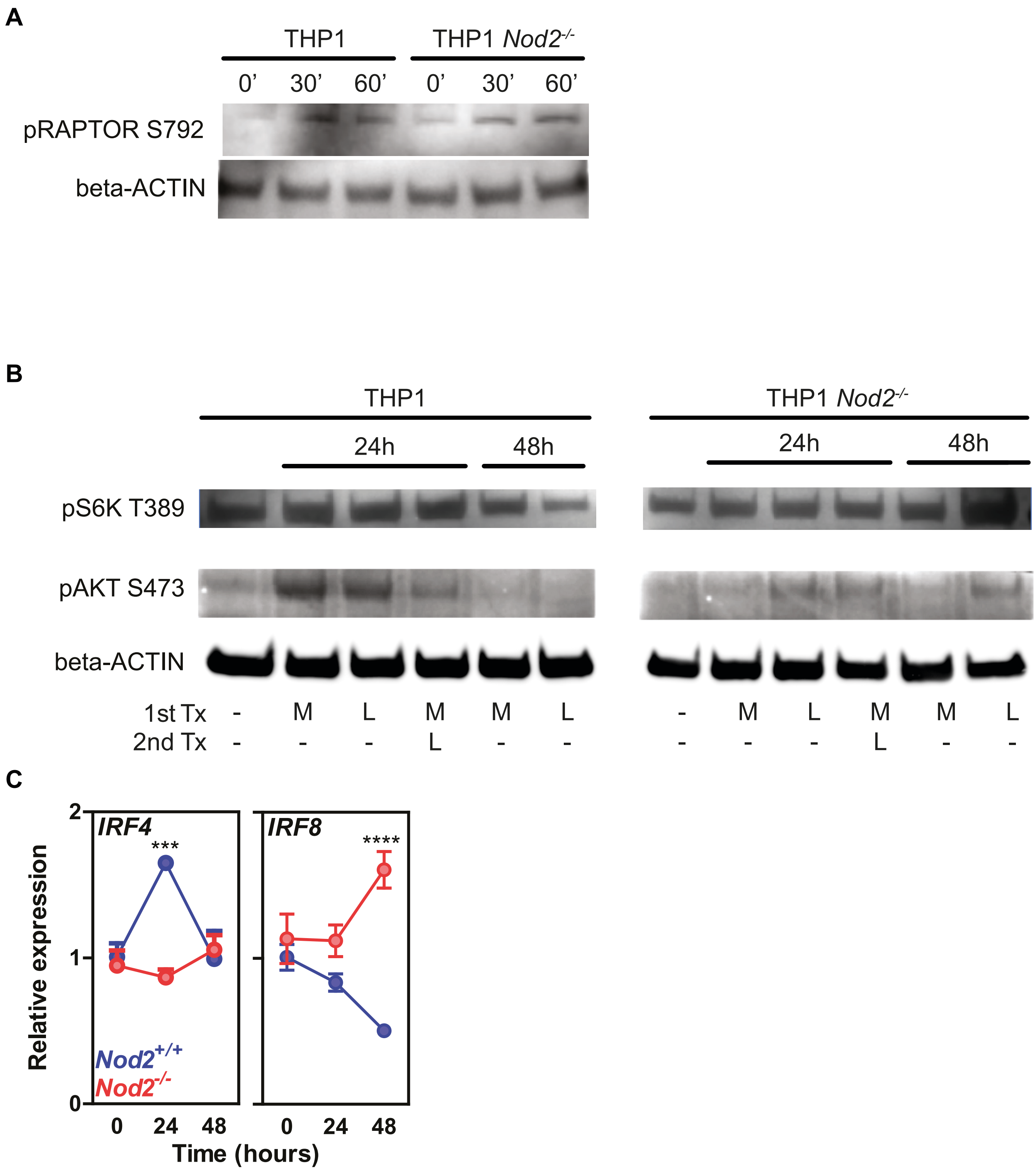
A
THP1
THP1 Nod2-/-
0' 30' 60' 0' 30' 60'
pRAPTOR S792
beta-ACTIN
B
THP1
THP1 Nod2-/-
24h
48h
pS6K T389
pAKT S473
beta-ACTIN
1st Tx - M L M M L - M L M M L
2nd Tx - - - L - - - - - L - -
C
IRF4
IRF8
***
****
Relative expression
Nod2+/+
Nod2-/-
0 24 48
Time (hours)

Supplementary Figure 9

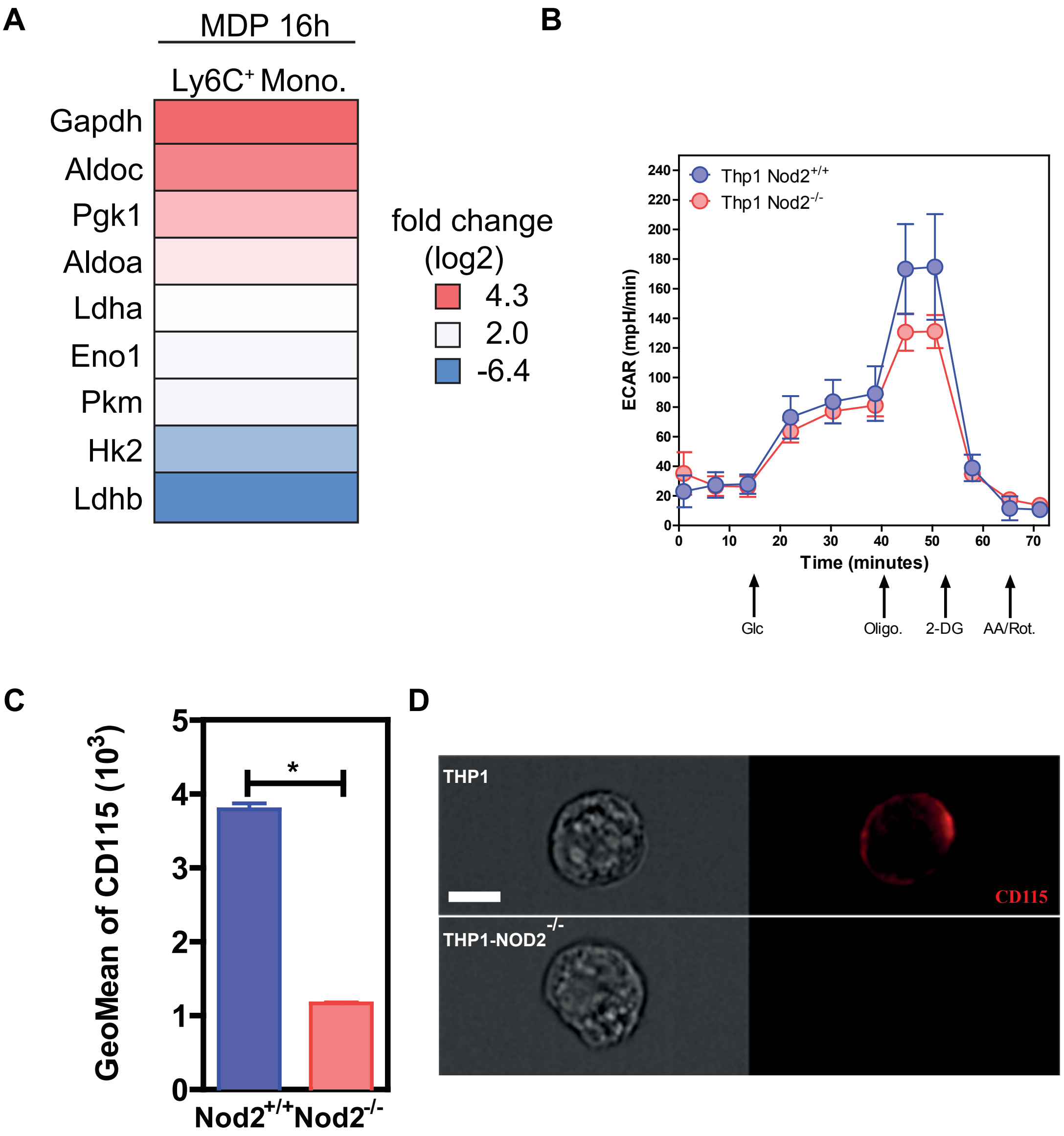

Supplementary Figure 10

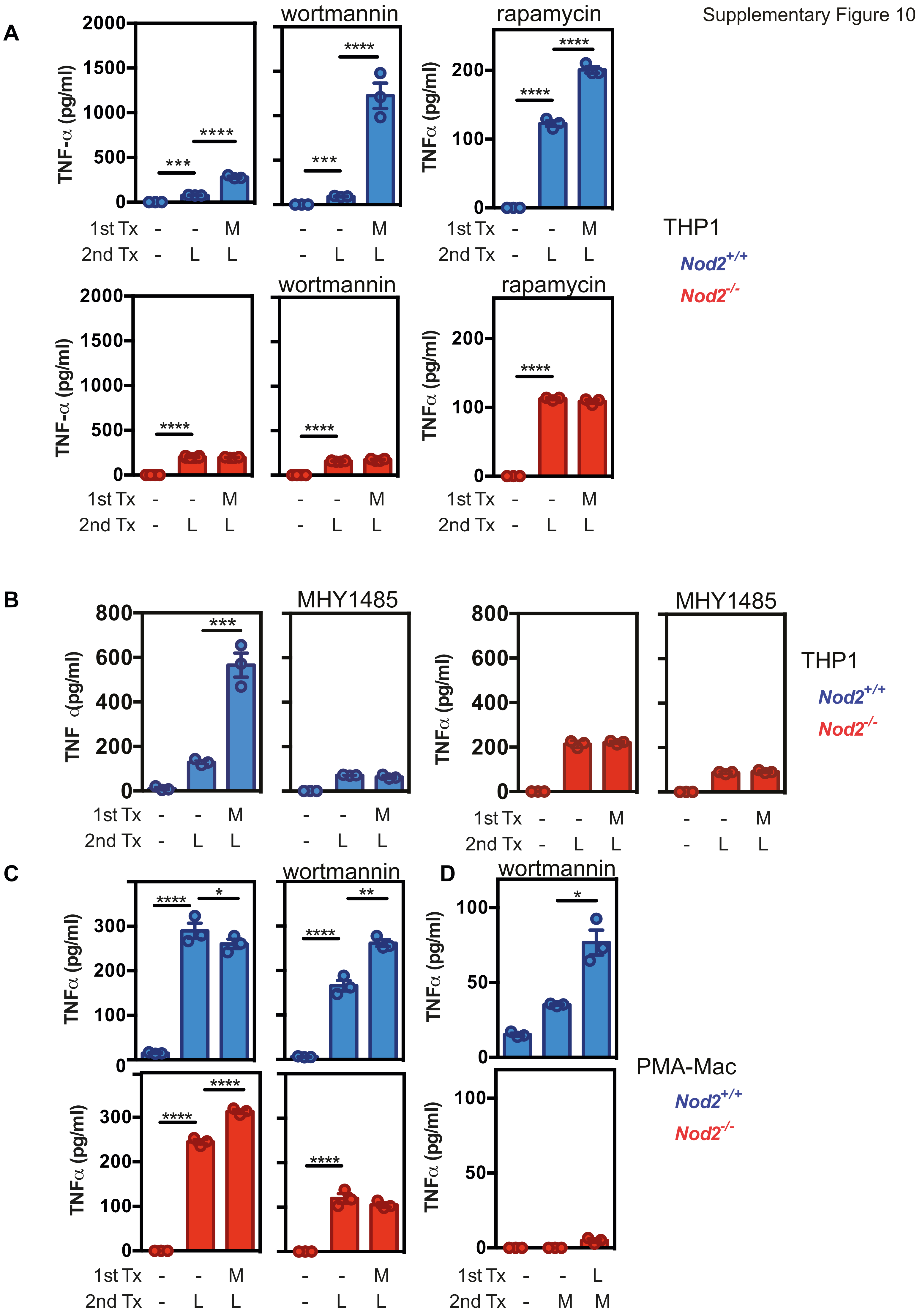

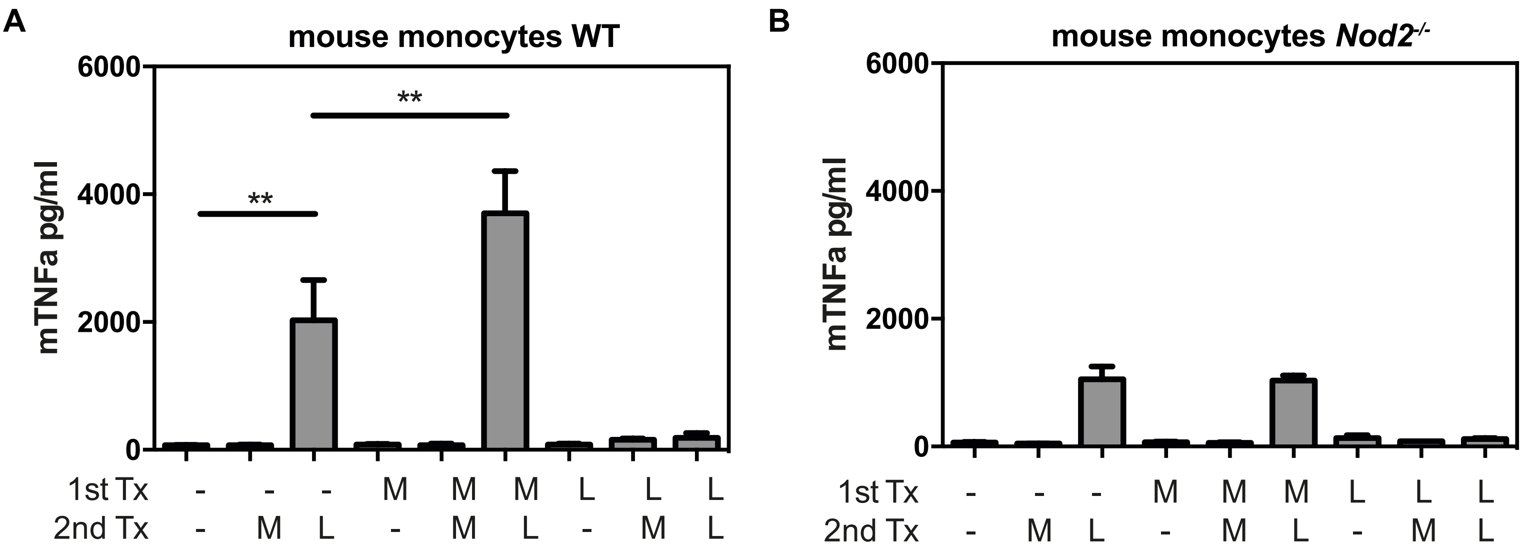
A
mouse monocytes WT
6000
4000
2000
0
mTNFa pg/ml
**
**
1st Tx - - - M M M L L L
2nd Tx - M L - M L - M L
B
mouse monocytes *Nod2*$^{-/-}$
6000
4000
2000
0
mTNFa pg/ml
1st Tx - - - M M M L L L
2nd Tx - M L - M L - M L

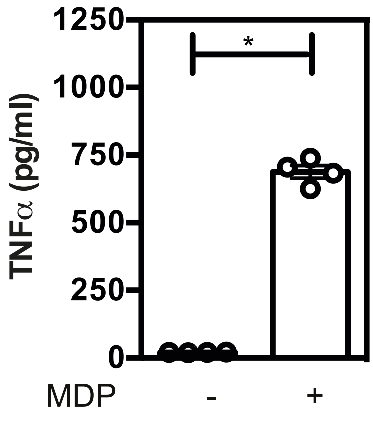
1250
1000
750
500
250
0
TNFα (pg/ml)
*
MDP
-
+
